# Areas of high risk for mammalian biodiversity and Nature’s Contributions to People under global warming

**DOI:** 10.1101/2024.08.30.610332

**Authors:** Marta Cimatti, Andrea Sacchi, Moreno Di Marco

## Abstract

Climate change has reached unprecedented levels, causing frequent extreme events like droughts and fires. Combined with land-use change, this crisis has impacted biodiversity, increasing species extinction rates, and Nature’s Contributions to People (NCP), degrading ecosystem functions. We developed a comparative extinction risk model for mammals sensitive to fire, drought, and extreme temperatures, utilizing a Random Forest algorithm to predict future extinction probabilities under different climatic scenarios. We then identified high-risk areas for both mammals and NCP under global warming, aiming to find synergies between biodiversity conservation and NCP preservation. Our results show that 288 out of 454 species (63%) face an increased extinction risk (mean increase 0.28), while 166 species (37%) show a predicted decrease (mean decrease 0.20) under the extreme "Fossil-fueled development" scenario. The highest risk increase was observed in Malaysia, Western Indonesia, Madagascar, Eastern Australia, and South Africa, under both pessimistic and optimistic ("Sustainability") scenarios. These regions also represent high-risk areas for several NCP: freshwater regulation, air quality, mitigation of extreme events. Preserving these high-risk regions is crucial for reducing habitat loss and human-induced extinctions. Safeguarding these ecosystems, which provide vital contributions like carbon storage, clean water, and extreme fire mitigation, should be a high priority. These regions warrant targeted policy and management interventions, including sustainable land-use practices and climate adaptation actions, to benefit both biodiversity and human well-being.

## Introduction

Human-induced climate change has now reached unprecedented levels, leading to more frequent and intense extreme weather events, such as floods, droughts, and fires which cause widespread adverse impacts to nature and people (Carnicer et al., 2022; Fischer & Knutti, 2015; He et al., 2019; IPCC, 2014, 2021, 2022; Ummenhofer & Meehl, 2017). These alterations have become a dominant threat to biodiversity (Bowman et al., 2020; Kelly et al., 2020; Maxwell et al., 2019; Ratnayake et al., 2019; Ward et al., 2020), affecting species distribution, phenology, population dynamics, and community structure (IPBES, 2019). The synergistic effect of heatwaves and drought can induce local wildlife declines and change habitat quality, resource availability, health, and welfare of wildlife (Maxwell et al., 2019; Sergio et al., 2018). For example, African elephant populations are sensitive to drought and to the extension of dry seasons, which impact their food and water availability leading to population reductions (Wato et al., 2016).

Climate change is also altering essential ecosystem functions, affecting water availability and quality, changing precipitation regimes and runoff (IPCC, 2021; Liu et al., 2022; Peña-Guerrero et al., 2020; Wang et al., 2020), or altering fire magnitude and frequency (Andela et al., 2017; Jolly et al., 2015; Rogers et al., 2020; Shlisky et al., 2009; Ward et al., 2020). Fire is a fundamental Earth system process, influencing vegetation distribution and ecosystem composition (Archibald et al., 2018; Roces-Díaz et al., 2022; Rogers et al., 2020) but more frequent high-severity fires can exceed the tolerance of species causing population reductions, local extinction and habitat destruction (Ward et al., 2020). Although a reduction in the global burned area has been recorded in recent decades, particularly in grasslands and savannahs (Andela et al., 2017), large fires are still documented in different parts of the globe, especially temperate and boreal zones, Mediterranean Europe, and southeastern Australia linked to abnormal temperatures and drought (Collins et al., 2022). Severe fire can negatively affect biodiversity by directly killing individuals, and by reducing the availability of food, water, and shelter for wildlife (Legge et al., 2021). For example, many Australian species already in a state of decline had a part of their habitat impacted by fires during 2019 and 2020, causing damage to the ecosystem structure which lead to a reduction in their capacity to recover (Ward et al., 2020, 2022).

In most ecosystems, changes in the number of species are the consequence of changes in major abiotic and disturbance factors, which are also intertwined in the generation of several regulating Nature’s Contributions to People (NCP) (Díaz et al., 2018, 2019). Since 1970, nature’s ability to support quality of life has declined for 14 of the 18 categories of NCP. These declines include all the regulating and non-material NCP, such as pollination, coastal resilience and soil protection, some of which are also projected to continue their decline in the future (Pereira et al., 2024). The impacts on ecosystems and NCP could be further enhanced by climate change, which is set to become an increasingly important factor into the future (IPBES 2019), by reducing the availability of areas with suitable climatic conditions for species, including areas that have not yet converted to human use (Di Marco et al. 2019; Arneth et al. 2020).

Hazards such as extreme weather, drought, or large and severe fire could also represent further risks for three important regulatory NCP: air quality regulation (NCP 3), regulation of freshwater quantity (NCP 6), and regulation of hazards and extreme events (NCP 9). Moreover, smoke released from large-scale fires could impact both human health through the decreased regulation of air quality (NCP 3) and wildlife health by influencing animal behaviour and decreasing air quality at a much larger spatial scale than the area burned (Sanderfoot et al., 2021; Vicente-Serrano et al., 2020; Xie et al., 2022). As consequence of the effects of global warming, both biodiversity and NCP will be impacted, generating high-risk areas for biodiversity and NCP which might lead to opportunities for synergistic conservation action.

The assessment of nature’s role in disaster risk reduction can be made by statistically comparing the impact of disasters in different areas and conducting comparative modelling analyses. A study by Chaplin-Kramer et al. (2019) exemplifies this approach, by evaluating coastal risk exposure through a ranked index that considers diverse physical factors such as sea level change and wave exposure. Consequently, vulnerability indices that account for specific natural characteristics can serve as a means to quantify nature’s impact on reducing hazards. The majority of NCP are provided through a combination of ecological interactions, biophysical process and human resources, consequently the methods used to evaluate the status of different NCP vary according to the study and NCP analysed. For example, regulatory NCP are not easy to measure because there are many abiotic factors involved. However, it is possible to estimate biophysical processes underlying the provision of NCP by measuring several indicators as suggested by the IPBES (Cimatti et al., 2023; IPBES, 2019; Jung et al., 2021; Y. Liu et al., 2023). Specifically, we measured the potential decrease of air quality regulation (NCP3) in terms of CO_2_ fire emission, the decrease of regulation of water quantity (NCP6), using water availability and actual evapotranspiration, and reduced hazard regulation (NCP9) by considering the extent of burned area.

Mammal species play a key role in numerous ecological functions, therefore their global decline can dramatically impact NCP provision (Hoffmann et al., 2011; Regan et al., 2015; Ripple et al., 2015). Mammals are involved in the provision of different regulating NCP such as pollination and seed dispersal (NCP2) or the regulation of detrimental organisms (NCP10), and provide important benefits to humans such as food, recreation and income (Material and non-material NCP) (Methorst et al., 2020; Rey et al., 2023). Mammals are also indirectly involved in fire control by reducing the fuel load through extensive grazing (Donaldson et al., 2018; Malhi et al., 2022). Understanding how the extinction risk of mammal species will change in response to global warming, and whether the same risk affects regulating NCP in the same regions, is fundamental to find synergies between the preservation of NCP and the conservation of biodiversity (Di Marco et al., 2016; Jung et al., 2021). Here we developed a comparative extinction risk model for mammal species sensitive to fire, drought, and extreme temperatures, and use it to predict their probability of extinction under future climate scenarios. We also projected risk to regulating NCP affected by the same stressors, and produced global maps to identify high-risk areas for biodiversity and NCP.

## Methods

We assessed the congruence between risk for mammalian biodiversity and risk for NCP provision under global warming (Fig. 1). We firstly developed a comparative extinction risk model considering both intrinsic and extrinsic variables for mammal species sensitive to fire, drought, and extreme temperatures according to the IUCN threats classification scheme. We used the result of this comparative extinction risk model to predict the probability of mammals to be threatened in the future. Secondly, we projected the risk of NCP loss affected by the same stressors (extrinsic variables) and produced global maps to identify areas of overlapping risk for biodiversity and NCP.

**Fig. 1:**
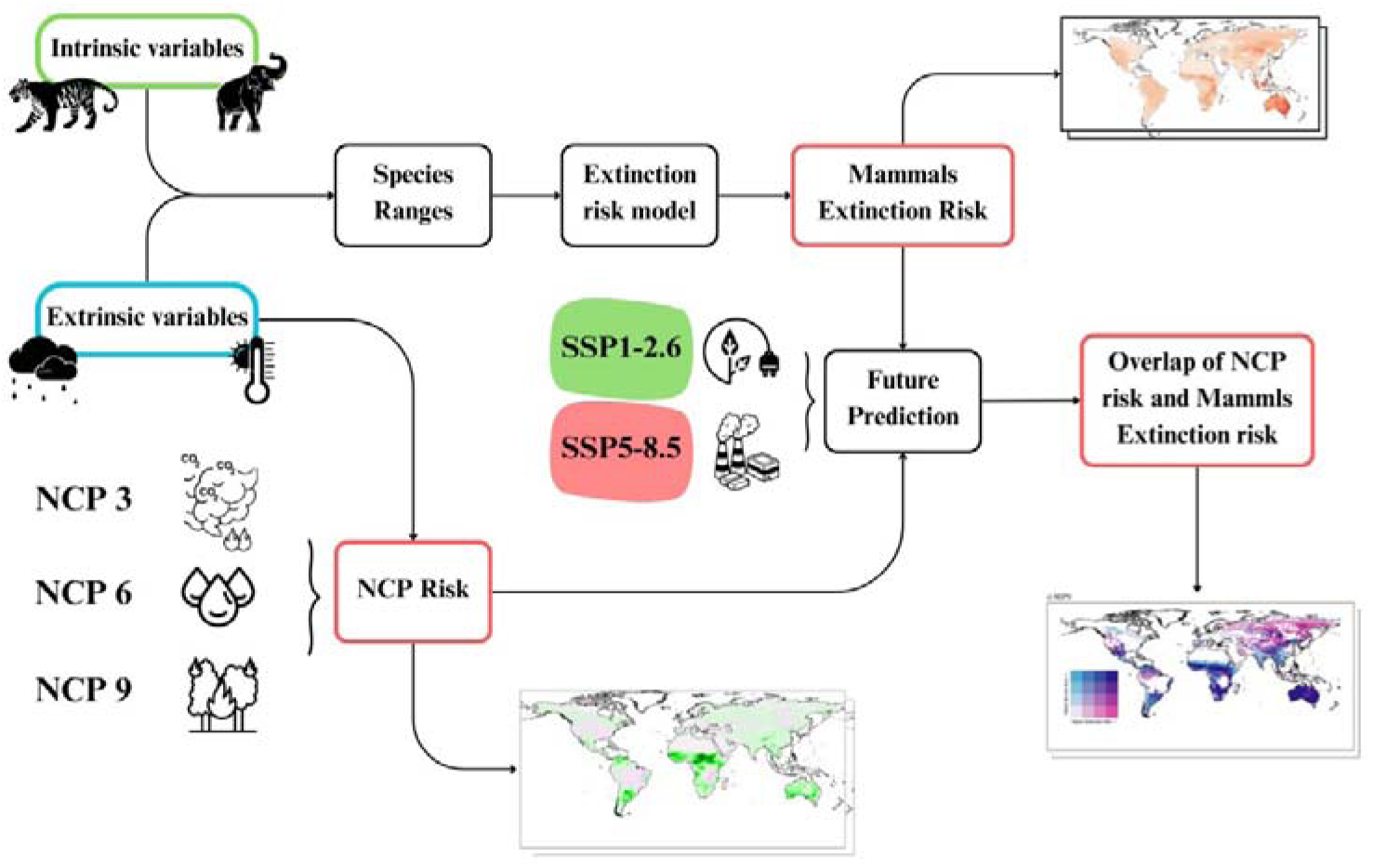
Methodological framework for selecting areas of overlapping risk for Biodiversity and Nature’s Contributions to People (NCP) provision under climate change.

### Species selection

We gathered mammal’s species distribution and extinction risk data from the IUCN Red List of Threatened Species (IUCN, 2022), selecting only those species classified as threatened by one or more among: increase in fire frequency/intensity, droughts, hot temperature extremes (Table S1) (IUCN, 2022; Salafsky et al., 2008). We excluded species listed as Extinct, Extinct in the Wild, or Data Deficient, resulting in 508 species being retained for further processing. We also filtered out from analyses those parts of the species ranges where species were introduced or had uncertain presence (Fig. S1).

### Threat data collection

We used intrinsic and extrinsic predictors to model the extinction risk for the selected species (Bielby et al., 2007; Di Marco et al., 2018; Pacifici et al., 2020; Pearson et al., 2014). We extracted life-history traits of mammal species from the Coalesced Mammal Database of Intrinsic and Extrinsic traits (COMBINE) (Soria et al., 2021). Following previous work (Davidson et al., 2017; Di Marco et al., 2018; Pacifici et al., 2020) we selected life-history variables that make species especially at risk such as body size, and speed of life history. Specifically, we considered two life-history traits linked with the “fast-slow continuum” concept, gestation length and interbirth interval, as surrogates for reproductive timing and reproductive output respectively (Bielby et al., 2007).

We obtained land use and environmental variables for two different time windows, a baseline period (1981-2010), and a future period (2041-2070) under different socio-economic development scenarios (SSP1-2.6 and SSP5-8.5). Scenario SSP1-2.6 “*Sustainability – Taking the green road”* is compliant with the Paris Agreement pursuing the effort of limiting warming to 2°C from pre-industrial levels, (Tebaldi et al., 2021; van Vuuren et al., 2011), while scenario SSP5-8.5 *“Fossil-fueled Development – Taking the Highway”* has a temperature increase projected between 4.1 and 4.7°C by 2100 (IPCC, 2014; Riahi et al., 2011). We retrieved land-use data from the LUH2 dataset available at http://luh.umd.edu/data.shtml (Hurtt et al., 2020), which provides maps at 0.25 x 0.25 degrees spatial resolution from year 850 to 2100. We chose to prioritize anthropogenic land-use classifications that are widely recognized for their negative impacts on biodiversity (Dragonetti et al., 2024; Newbold et al., 2015). Thus we aggregated the different classes of anthropogenic land-use into one macro-class called anthropic, multiplying them with the grid-cell area. We resampled each layer at 1 km resolution using bilinear interpolation, to extract average land-use conditions from within each species range without losing marginal portions of the range due to the coarse native resolution of land-use data.

We then retrieved several variables which represent potential environmental condition associated with the threats considered in our analysis (Table 1and Table S2) as a proxy of NCP indicators (Chaplin-Kramer et al., 2019; Jung et al., 2021; Y. Liu et al., 2023, Shuai et al., 2021; Visconti et al., 2016; Ward et al., 2020): temperature, water availability, actual evapotranspiration, burned area, and CO_2_ emissions from fire.

**Table 1:** List of life-history traits and environmental variables used as predictors of extinction risk in the random forest model with VIF < 4 (full variable list in Table S2). Data sources are cited and described in the main text.

| Variable name | Definition | Summary statistics | Unit of measures | Data source |
| --- | --- | --- | --- | --- |
| Adult mass | Body mass of an adult individual in grams | Mean: 29.238<br>Range: 0.002 to 3220 | Kg | COMBINE (Soria et al., 2021) |
| Gestation length | Length of time of foetal growth in days | Mean: 98.11<br>Range: 12 to 642.84 | days | COMBINE (Soria et al., 2021) |
| Interbirth interval | Time between reproduction events in days | Mean: 291.41<br>Range: 23.71 to 1709.58 | days | COMBINE (Soria et al., 2021) |
| Order | Mammals order |  | categorical | COMBINE (Soria et al., 2021) |
| Minimum Water Availability | Minimum annual water availability | Mean: -27.35<br>Range: -179.58 to 399.66 | mm/month | CHELSA /CMIP6<br>(https://chelsa-climate.org/downloads/)<br>(https://esgf-node.llnl.gov/search/cmip6/) |
| Minimum Burned Area | Minimum annual burned area | Mean: 0.04<br>Range: 0 to 0.29 | Area (%) | CMIP6<br>(https://esgf-node.llnl.gov/search/cmip6/) |
| Maximum CO2 Emission | Maximum annual CO2 Emission from fire | Mean: 0.007<br>Range: 0 to 0.03 | kg/m<sup>2</sup> | CMIP6<br>(https://esgf-node.llnl.gov/search/cmip6/) |
| Maximum Actual Evapotranspiration | Maximum annual actual evapotranspiration | Mean: 100.86<br>Range: 4.04 to 255.03 | mm/month | CMIP6<br>(https://esgf-node.llnl.gov/search/cmip6/) |
| Anthropic | Sum of Cropland (C3 and C4 types), Pastureland and Urban | Mean: 116.13<br>Range: 0 to 512.87 | km<sup>2</sup> | LUH (Hurt et al., 2020) |
| Warm days | Percentage of days with maximum temperature above the corresponding calendar day 90th percentile of maximum temperature for a 5-day moving window in the base period | Mean: 13.70<br>Range: 9.81 to 27.68 | % | ETCCDI<br>(Kim et al., 2020) |
| Minimum value of daily maximum temperature | Minimum of daily maximum temperature in period (year or month, TXn) | Mean: 14.16<br>Range: -40.79 to 25.65 | °C | ETCCDI<br>(Kim et al., 2020) |
| Warm spell duration index | Annual count of days with at least 6 consecutive days when daily maximum temperature is above the calendar day 90th percentile of maximum temperature centered on a 5-day sliding window during the base period (wsdi) | Mean: 12.94<br>Range: 0.77 to 28.19 | days | ETCCDI<br>(Kim et al., 2020) |
| Extreme fire weather | Number of days with extreme fire weather (fwixd) | Mean: 19.71<br>Range: 12.69 to 36.01 | days | FWI (Quilcaille et al., 2023) |
| Length of the fire season | length of the fire season (fwils) | Mean: 14.25<br>Range: 10.59 to 18.55 | days | FWI (Quilcaille et al., 2023) |

We collected temperature and precipitation data from the Chelsa dataset at 1km res (https://chelsa-climate.org/downloads/) (Karger et al., 2017); data of burned area, CO_2_ from fire emissions, and actual evapotranspiration were retrieved from the CMIP6 dataset (https://esgf-node.llnl.gov/search/cmip6/), using the R package “epwshiftr” (Jia & Chong, 2021) (Table S2).

Water availability (WA) was calculated as the net difference between precipitation (P) and actual evapotranspiration (AET) (Konapala et al., 2020) as [mm/month] [1]:

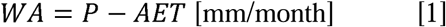

Finally, we included the actual evapotranspiration (AET) as a drought indicator. AET is defined as the actual amount of water removed from a surface area through a combination of evaporation from soil and plant surfaces, and transpiration through plant canopies. Due to the warming climate, AET is expected to increase in the future, likely leading to more frequent and intense extreme events (Nooni et al., 2021).

We collected a set of fire weather index (FWI) variables from Quilcalle et al (Quilcaille et al., 2023), namely length of the fire season (fwils), and the number of days with extreme fire weather (fwixd), defined in Abatzoglou et al. (Abatzoglou et al., 2019) and Jolly et al. (Jolly et al., 2015) as the conditions favouring the ignition and persistence of fires, characterized by the convergence of hot, dry, and windy events.

Finally, in order to include the effects of extreme weather events, we collected data from Copernicus Climate Change Service, selecting climate extreme indices respectively related to hot extreme temperature and dry condition (Table1, Table S2) as defined by the Expert Team on Climate Change Detection and Indices (ETCCDI) (Kim et al., 2020).

For most of the above-listed environmental variables we selected five priority climate models (General Circulation Models, GCMs) following guidelines from the Inter-Sectoral Impact Model Intercomparison Project Phase 3 (ISIMIP) (Frieler et al., 2024; Lange, 2019). For the burned area and fire emission the top priority GCMs were not available, thus we used all the other GCMs present on the CMIP6 repository (Table S3). We then averaged the values across all GCMs for each climate scenario. Finally, we extracted the median of the maximum, minimum and average values of each variable (for FWI and ETCCDI only the median of average values) (Table S2) within the range of each species for the analyzed periods using the packages *raster, sf, extactextract* (Baston, 2023; Hijmans, 2016; Pebesma, 2016) of the R software (version 4.3.1). Finally, we excluded 44 species, for which some variable values were missing (e.g. island species, for which original data resolution was too coarse), resulting in 454 species being analyzed.

### Extinction risk modeling

We built an extinction risk model to predict the probability of species being threatened with extinction based on their intrinsic life-history traits and the value of extrinsic variables within their distributions. We classified species as “threatened” if they were assessed as Vulnerable, Endangered, or Critically Endangered in the IUCN Red List, and “non threatened” otherwise. We used the *ranger* package (Wright & Ziegler, 2017) implemented in R to run a Random Forest classification model based on the intrinsic and extrinsic variables selected (Table 1). We adopted a 10-fold cross validation, simultaneously tuning the hyperparameters *mtry*, *min.node size* and *sample size* using a grid search. Furthermore, in order to reduce collinearity in the model, we tested different combination of variables given by variance inflation factors VIF threshold between 10 and 3 (Alin, 2010; Elith et al., 2012; Zuur et al., 2010), finally choosing the VIF threshold and hyperparameters combination that resulted in a higher TSS.

The Random Forest model produced continuous estimates of the probability for each species to be threatened, and in order to validate model’s performance we binarized that probability (threatened vs non-threatened) considering species with predicted values of probability of being threatened higher than 0.5 as Threatened. We measured the model accuracy using the TSS which ranges between -1 and 1, where +1 indicates perfect agreement, and values of zero or less indicate a performance no better than random (Allouche et al., 2006). Furthermore, to investigate random forest model predictive ability, we measured TSS also through a taxonomic block validation, in which all species in a taxonomic family were left out of the model training and then used for validating model’s predictions, iteratively across all families with at least 15 species. While random cross validation is a good indication of model’s ability to interpolate data, block validation is an indication of model’s ability to extrapolate predictions to new (independent) data (Di Marco, 2022).

We used the Random Forest classification model to predict the probability of species to be threatened in the present and under two scenarios of future environmental change: a low emission scenario SSP1-2.6, and a very high SSP5-8.5. For generating future probabilities, we recalculated the value of extrinsic variables associated to climatic conditions for each scenario and re-predicted each species risk using the same model trained on present-day variables. Intrinsic variables, i.e. biological traits, were retained as static through time.

In addition, we determined the predictor importance using the Mean Decrease Accuracy, which quantifies the importance of a variable by measuring the change in prediction accuracy when the values of the variable are randomly permuted compared to the original observations (Calle & Urrea, 2011). We also represented the relationship between the predictors and the response variable by constructing partial dependence plots for each independent variable with the package *pdp* (Friedman, 2001; Greenwell, 2017). We applied a locally weighted least square regression (loess) to fit a smooth curve on the partial dependence plot to reduce the effect of overfitting in the model.

Finally, for each species we calculated the difference in probability of being threatened (*Δp*) between the future (*p_f_*) and present (*p_h_*) under each scenario [3].

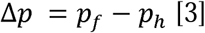

A positive Δ*p* characterized species that had an increase in their probability of being threatened, and a negative Δ*p* characterized species that had a reduction in their probability of being threatened.

### Identification of high-risk areas overlap between biodiversity and NCP

We created a cumulative map of extinction risk for each scenario, by summing all the Δ*p* values of species in each grid cell location of a raster map. We produced separate maps for species facing an increase in extinction risk (“high-risk areas” map) or a decrease in risk (“low-risk areas” map).

At the same time, we delineated the spatial distribution of areas that are predicted to suffer a loss of regulating NCP, developing different maps under the two scenarios (SSP1-2.6 and SSP5-8.5). We mapped change in freshwater quantity (NCP 6) by measuring minimum water availability, change in extreme events (NCP 9) by measuring the fraction of burned area, and the change in air quality (NCP 3) by mapping CO_2_ emissions from fire. Finally, we created bivariate maps of species vs NCP to identify high-risk areas for both.

## Results

### Extinction risk prediction

The Random Forest model for predicting extinction risk had good classification accuracy, with 78% species correctly classified. The model had high sensitivity (68% correctly classified threatened species), and very high specificity (87% correctly classified non-threatened species), resulting in a TSS value of 0.55 (on a scale between -1 and 1). Overall, 195 species were predicted as threatened and 259 species as non-threatened by the model, using a probability cutoff of 0.5. During taxonomic block validation, model’s performance was lower with a mean TSS of 0.25 but there were large differences among families; the model had the highest predictive performance (TSS of 0.61) for Soricidae and lowest performance for Dasyuridae and Bovidae (TSS of 0 and -0.17 respectively; Table S4).

We calculated the importance of each variable according to the decrease in classification accuracy and found overall a higher predictive importance for intrinsic (life-traits) compared to extrinsic (climatic/land use) variables (Fig. S2). Adult body mass was the most important variable, followed by other known correlates of risk such as gestation length interbirth interval. The probability of being threatened was positively correlated with body mass and reproductive traits, confirming large-bodied slow-reproducing species face higher risk (Fig. 2).

**Fig. 2:**
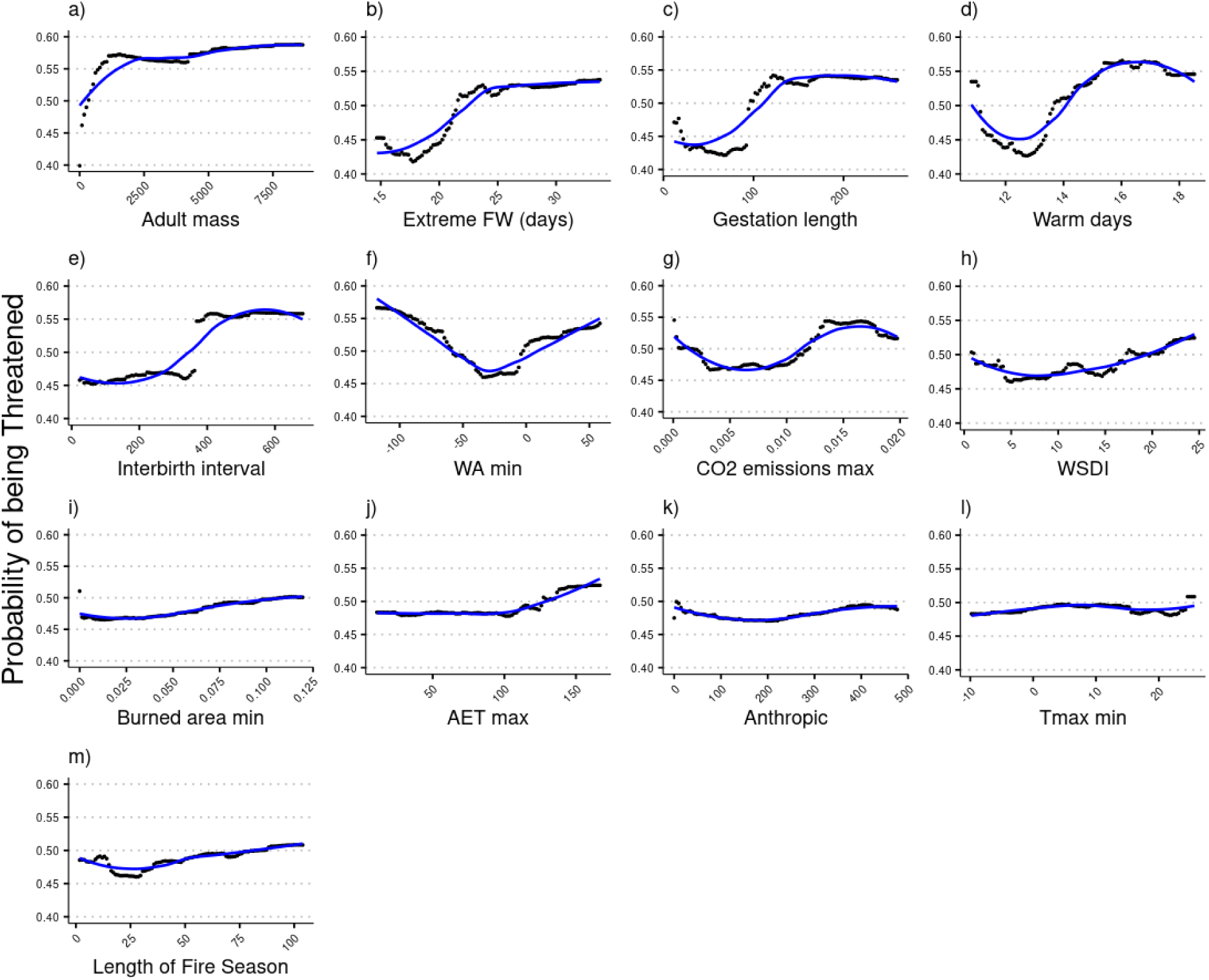
Partial dependence plots of the most important model variables and those used as NCP indicator (Burned area, Water availability, and CO2 emissions from fire), showing the effect of predictors on the probability of extinction. a) Adult mass, b) Extreme fire weather days, c) Gestation length, d) Warm days, e)Interbirth interval, f) Minimum water availability, g) Maximum CO2 emission from fire, h) Warm Spell Duration Index, i) Minimum burned area, j) Maximum actual evapotranspiration, k) Anthropic land use, l) Minimum of daily maximum temperature m) length of Fire Season. Variables are ordered according to their mean ranked importance for the Mean Decreased Accuracy.

The most important extrinsic variables were extreme fire weather days, the percentage of warm days and the followed by minimum water availability and minimum burned area. The probability of being threatened correlated positively with the extreme fire weather and the percentage of warm days while the response of minimum water availability was idiosyncratic. The correlation with the probability of being threatened was positive also for the CO_2_ emission from fires and the minimum burned area and the rest of extrinsic variables.

#### 1.1.2. Scenarios of future risk

When using the random forest model to predict the future risk of species being threatened, we found very similar results across scenarios, as a consequence of life-history traits (i.e. static through time and between scenarios) being the most important predictors in the model (Fig. S2). Under the historical period, the median probability of being threatened across the 454 modelled species was 0.46, while this increased in the future to 0.567 and 0.569 respectively under scenarios SSP1-2.6 and SSP5-8.5. The probability of being threatened increased for 287 species and decreased for 167 species under scenario SSP1-2.6. While for scenario SSP5-8.5, the probability of being threatened increased for 288 species and decreased for 166 (Fig. S3-S4).

Taxonomic orders facing higher risk under scenario SSP5-8.5 included Notoryctemorphia, Peramelemorphia and Eulipotyphla (Fig. S5-S7). Species currently in the category Least Concern and Near Threatened faced the highest increase in extinction risk, with 79% and 77% of species belonging to those classes respectively increasing their risk under the scenario SSP5-8.5 (Fig. S8).

### Global hotspots of extinction risk and NCP risk from climate change

We mapped the aggregated change in extinction risk for species between the present and the future and found only localized differences among realms and scenarios in terms of both positive and negative change in risk (Fig. 3). For example, the highest positive change values were observed on Thailand and Malaysia, Borneo and in Madagascar both under scenarios SSP1-2.6 and SSP5-8.5 followed by the East coast of Australia, Gobi Desert and Mongolian-Manchurian steppe (Fig.3, Fig. S9).

**Fig. 3:**
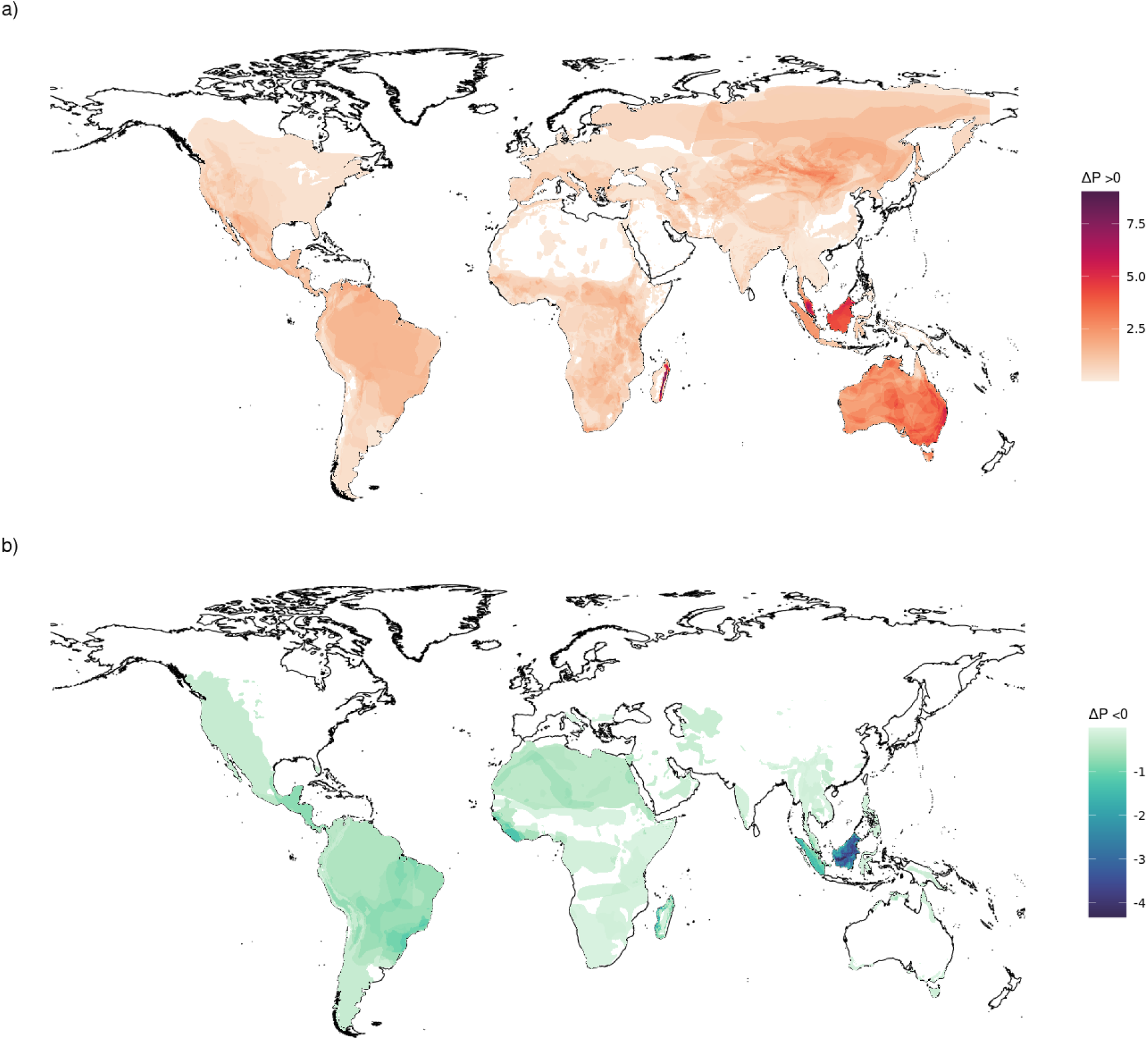
Global cumulative change (Δp) in species extinction risk for species showing a) a risk increase and b) a risk decrease, under scenario SSP5-8.5.

Also, the global maps produced showed similarities among scenarios in terms of change in the probability of being threatened. For example, in the Neotropical realm, the area with high negative change in extinction risk was the in the central zone of South America, in particular most of the Bolivian and north Chilean Andes, Parà and Sao Paolo Brazilian states, while in the Nearctic realm the Rocky Mountains had a high negative change in probability of extinction. In the eastern Palearctic realm, regions near Caspian Sea were also characterized by species with a decrease in extinction risk compared to the present time window. Surprisingly, also parts of the West African coast and parts of Southeast Asia, in particular the Indo-Malay realm were characterized by the highest increase in extinction risk and hosted numerous species with a negative change in extinction risk, for the most part belonging to Primates order.

When mapping the change in the provision of regulating NCP, we found the maximum CO_2_ fire emissions, an indicator of air quality regulation (NCP 3) showed an increase in few areas in north America, east Amazon, Uruguay and Patagonia, and a strong increase in Cameroon, the Central African Republic, Northwestern Russia and Ural region, Tibet and South-east Australia in scenario SSP5-8.5 compared to the baseline period (Fig. S10). The minimum water availability instead was projected to decrease under both scenarios, which indicates a future reduction of the provision of water quantity regulation (NCP 6), with particularly low values in the Amazon, the north American east coast, South Africa, parts of Europe, Southeast Asia, and Australia (Fig. S11). Moreover, we observed an increase in the minimum burned area and in the actual evapotranspiration (NCP 9), under both scenarios (Fig. S12-S13). The minimum burned area increased especially under the most pessimistic scenario SSP5-8.5 in Venezuela, Colombia, Uruguay, Congo, Southern Africa, and large parts of Asia and Australia (Fig. S12) while the two most important extrinsic variable, number of days with extreme fire weather and percentage of warm days, increase overall across the globe (Fig. S14-S15).

We produced bivariate maps of the relationship between NCP risk and species extinction risk (Fig. 4, Fig. S17). Under scenario SSP5-8.5 there were different areas where an increase in CO_2_ from fire emissions (decrease in air quality provision) overlapped with areas characterized by an increase in mammals’ probability of extinction, such as the great lakes and great plains regions of North America, Colombia, Perú, part of the Amazon, and parts of Argentina, tropical rain forests of Central Africa, a small spot in eastern Europe, and large part of Ural and Siberia region in Russia, Tibetan plateau and Sichuan region, and south Australia. The overlap between the decrease in minimum water availability and mammals’ extinction risk had a wider distribution with high value of extinction risk corresponding to a decrease in freshwater quantity regulation also in Sub-Saharan Africa, Madagascar, the majority of North and South America, and parts of Southeast Asia and Australia. Scenario SSP5-8.5 showed high increases in both extinction risk and burned area extent in the Neotropical, Afrotropical, and Palearctic realms. In particular, the southern part of the Rocky Mountains, northern part of Colombia and Venezuela, Caatinga regions and Pampas will be affected by an increase in burned area simultaneous to an increase in mammal extinction risk. Large parts of southern Europe will host high-risk areas of fire and species risk, together with Southern Africa, from Tibetan plateau through parts of the Mongolian-Manchurian steppe to Siberia, and almost the whole of Australasia (Fig. 4).

**Fig. 4:**
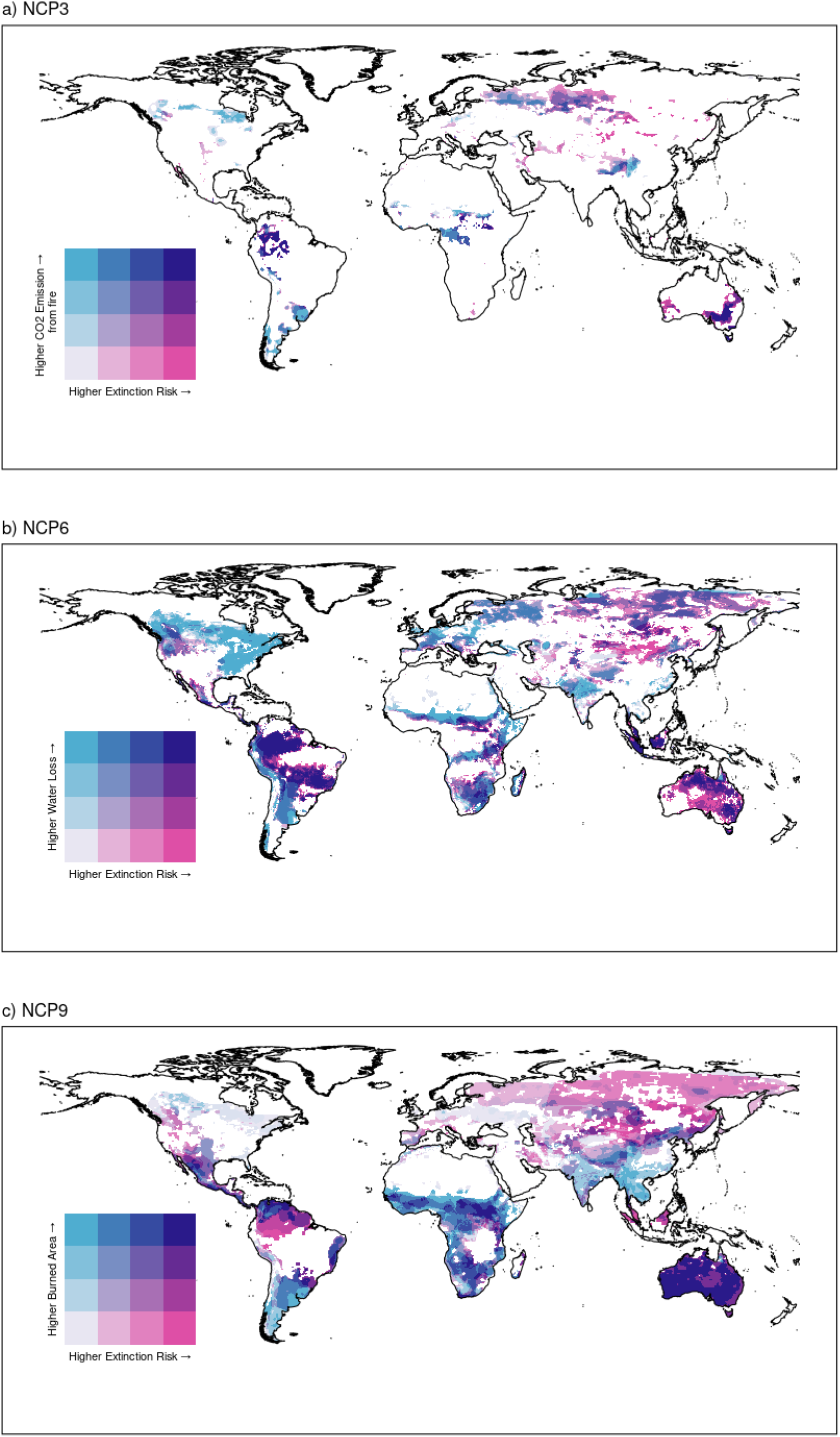
Global bivariate maps indicating high-risk areas of high cumulative risk for species and high risk of NCP loss: a) CO_2_ fire emission, indicator of Air quality regulation (NCP 3), b) Water availability indicator of freshwater quantity regulation (NCP 6), and c) Burned area, indicator of regulation of extreme event (NCP 9). The map reports relative increase in extinction risk for species expected to have higher risk in the future and increase the proxy for NCP under SSP5-8.5 scenario. Hotpots of species and NCP risk are identified in purple.

## Discussion

The increased frequency and intensity of ecosystem disturbance associated with global warming has already been investigated for mammals (Ameca y Juárez et al., 2013; Isaac, 2009; Ward et al., 2020), and here we predict how future warming levels will affect extinction risk for species classified as sensitive to droughts, fire, and high temperature. Risk was especially associated with life-history traits such as body mass and gestation length, supporting results of previous studies (Cardillo et al., 2008; Davidson et al., 2017; Lee & Jetz, 2011). Large body mass has been linked to elevated extinction risk because large mammals have lower reproductive rates, slower population growth (Cardillo et al., 2008), and have experienced large range contractions of their range from human activities (Pacifici et al., 2020). We found extrinsic variables were of secondary importance compared to the above-mentioned life-history traits (Fig. S2). This result partly contrasts with that of recent studies on mammals (Di Marco et al., 2018; Murray et al., 2014) where extrinsic variables (i.e. pressure level) played a dominant role. This is likely related to the particular subset of species analysed here, which do not represent the overall threat status of mammals but rather a selection of species particularly sensitive to environmental change according to the IUCN Red List assessors. Furthermore, secondary importance of extrinsic variables could be as a consequence of the species threats defined by IUCN experts describe the environmental variables to which the species are overall sensitive to, but they do not directly measure the exposure, which is instead related to their threat status.

While extrinsic variables were less significant in the model, they still played a crucial role in influencing extinction risk. Specifically, we found that the percentage of warm days, burned area and extreme fire weather days had a clear positive correlation with the probability of extinction while water availability had an idiosyncratic response, with very low or very high values of water availability correlated to a higher probability of being threatened. This counterintuitive response could be due to spurious correlation linked to the spatial distribution of mammal species at risk that we analysed. Indeed, our dataset, included several threatened species distributed in areas that are predicted to become drier, such as the Amazon, but also species in areas with high values of water availability, especially in Southeast Asia.

Our results of the probability of being threatened does not differ significantly from results presented in previous studies (Cardillo et al., 2006, 2023; Davidson et al., 2017; Pacifici et al., 2018; Torres-Romero et al., 2024). For example, we found Madagascar will be a high-risk area for mammals, especially along the moist lowland forest in the eastern part of the island. Madagascar has been identified as a conservation hotspot in several studies due to its high level of endemic and small-ranged species (Jenkins et al., 2013; Mittermeier et al., 2004; Myers, 2003; Myers et al., 2000), and the presence of several threats linked to human pressure, including the fragmentation and loss of pristine forest (Harper et al., 2007; Vieilledent et al., 2018). We found a decrease in CO_2_ fire emissions and burned area in this region reflecting past trends (Phelps et al., 2022), nevertheless, the slash and-burn approach employed to clear forests for agriculture and to rejuvenate cattle pastures remains a significant cause of mortality for numerous Malagasy forest species (Frappier-Brinton & Lehman, 2022; Kull & Laris, 2009). We also found that the percentage of warm days and the number extreme fire weather days are projected to increase in the future in Madagascar, especially under scenario SSP5-8.5 (Fig. S14-S15). This confirms that Madagascar is concurrently highly vulnerable to climate change and natural catastrophe including cyclones, flooding and droughts and it is expected that these events will further intensify in the future (Tadross et al., 2008; Tanteliniaina et al., 2021).

Southeast Asia is another high-risk area that emerged from our analysis. This area is considered one of the most biodiverse globally (Myers et al., 2000), but nowadays faces some of the highest human pressure from hunting deforestation, and fire (Chiaverini et al., 2022; Hughes, 2017; Sodhi et al., 2004). We found additional risk might be posed by future climate change, as global projections show increases in the number of warm days and extreme fire weather days (Fig. S14-15) and increase in actual evapotranspiration potentially leading to droughts in a region which is already facing water stress (Hughes, 2017; Lickley & Solomon, 2018; Zhang et al., 2016). Hotter and drier environments could affect mammal’s water balance leading to a reduction in the fitness, survivability and reproductive success of species and consequently to an increase of the probability of extinction (Fuller et al., 2016; Milligan & Lloyd, 2009). Thus, drought and water scarcity are significant drivers of extinction risk in Southeast Asia, both currently and in the future. Projections, particularly under the SSP5-8.5 scenario, indicate an increase in mean drought characteristics, such as its duration, intensity, and severity further amplifying this threat (Supharatid & Nafung, 2021).

Our model also predicted a decrease in extinction risk for several species, highlighting the presence of “low-risk” areas, similarly to results found in Allan et al 2019 (Allan et al., 2019). Our and Allan et al., results do not exactly overlap, since we use a limited set of species and different threats but we also highlight areas with a decrease in extinction risk in South and Northwest America, the African east coast (i.e. in Liberia, in this case exactly as in Allan et al.), and Borneo and Sumatra Islands. The little difference in the spatial distribution of high-risk areas and low-risk areas between climate change scenarios in Indo-Malay and Afrotropic realms is based on species diversity and the varying sensitivity of individual species to threats. Coincidently, we found an high concentration of Primate species, which were the order that had the largest reduction in probability of being threatened (Fig. S7-S10), probably because they are concentrated in areas characterized by a predicted decrease in burned area and related emissions. The spatial variation in the pressures faced by the different species could explain the reduction in probability of being threatened. For example, our results highlight an increase in water availability in some parts of South America such as Venezuela, Rio Grande Do Sul in Brazil, some countries facing Guinean Gulf, but also New Guinea under scenario SSP5-8.5, while parts of Southeast Asia are characterized by a decrease burned area and fire CO2 emission. These patterns could also be attributed to the high importance of intrinsic variables, that do not change with future projections, compared to extrinsic variables. Furthermore, considering also that threat risk is shaped by interactions that arise between species’ intrinsic traits and the extrinsic factors (Duncan et al., 2012; Murray et al., 2014), most of the species that face a decrease in extinction risk have also a small body size, which is correlated to a faster ability to adapt to rapid environmental changes (Fig. S19) (Pacifici et al., 2018).

We identified several areas where high cumulative species risk coincides with a high risk of NCP loss, despite the greater significance of intrinsic variables compared to extrinsic ones in the extinction risk model. Regions of shared risk are those facing future increases in hot and dry climate conditions, such as the Amazon, Mediterranean, southern Africa, parts of Australia and the Southeast Asian monsoon region (Cook et al., 2020; Liu et al., 2022; Richardson et al., 2022). The corresponding decrease in water availability in these areas results in reduced water quantity regulation (NCP 6), with climate change further aggravating drought by either enhancing evapotranspiration or suppressing precipitation (Liu et al., 2022; Zhang et al., 2021).

Our results reflects the projected increase in fire intensity and extent from accelerated high-latitude warming and tropical and subtropical drying predicted by CMIP6 models, in line with other studies (Abatzoglou et al., 2020; Quilcaille et al., 2023; Seneviratne et al., 2021). One of the regions with the largest increases in burned area and CO_2_ fire emissions globally is the western United States. The aggressive suppression of systems adapted to frequent low-severity fires, and the expansion of the wildland urban interface, have created conditions for increasingly dangerous fires in this region (Rogers et al., 2020). The impact of wildfire on air pollution are likely to become more severe due to climate change, especially under more extreme scenarios (Wu et al., 2021; Yang et al., 2022). Fire emissions can considerably impact public health on large scales (Brey et al., 2020) by reducing air quality over extensive areas due to the sizable amounts of particles and trace of harmful gases present in fire emissions (Van Der Werf et al., 2017; Yang et al., 2022).

The two largest biomes of Brazil, the Amazon and the Cerrado, will also represent areas of high fire risk and extinction risk in the future. Fires in the Amazon rainforest typically indicate deforestation and are usually started by humans for agriculture or land clearing (Soares-Filho et al., 2012).

By contrast, the fires occurring in the Cerrado, a vast savanna, are primarily natural due to the prevalence of fire-prone vegetation. However, despite the biome fire-prone nature, an increase in the frequency and intensity of drought induced by climate change could disrupt these ecosystems, making recovery from previous wildfire unfeasible for fauna and flora (Li et al., 2021). High-risk areas for increased CO_2_ emissions and burned area did not always match. Zheng et al. (Zheng et al., 2021) already highlighted this pattern for CO_2_ emissions from 2000 to 2019, proposing that despite global burned areas having declined, the CO_2_ emissions from global fires did not diminish. This is due to trees having a greater fuel load and releasing more carbon into the atmosphere per unit of area burned compared to grasses. Burning in increasingly fragmented forest may initiate a self-reinforcing loop, which augments the rainforest fragmentation and affects its carbon storage, transforming carbon sink forest into a carbon source (Driscoll et al., 2021).

Exploring and including in our analysis the relationship and feedback between fire and mammal biodiversity would be of high interest for biodiversity conservation under global change. Nonetheless, understanding and defining how mammals can impact and mitigate the effect of fires is based on a variety of characteristics such as population density, feeding strategy (grazer or browser) and foraging sources (grass or leaves) (Fairfax & Whittle, 2020; Foster et al., 2020; Young et al., 2022). There are different studies at local scales that try to model how fire and mammal herbivory, two different plant “consumers”, control the fuel load and vegetation structure in many terrestrial ecosystems (Donaldson et al., 2018; Fairfax & Whittle, 2020; Zylinski et al., 2022) but large-scale data (such as those needed here) are not yet available (e.g. present and future species abundance, plant diet preferences, rate of vegetation consumption).

Finally, we found several areas where high cumulative risk for species matches a risk of NCP loss from global warming under different scenarios, even if limiting the global warming to below 2°C, in accordance with the Paris Agreement (represented here by the scenario SSP1-2.6). Cumulative risk will be especially high in the developing tropical regions, that will face extensive land use change due to the increased demand for cropland, logging and households, but our results also highlighted areas outside tropical region that will be exposed to higher fire and drought risk, such as large spots in North America and Patagonia parts of Europe and Siberia. Areas with large overlaps of extinction risk and NCP risk should be considered as priority for interventions, as conservation action would serve a dual purpose. Savannas and tropical forests are especially at risk of facing biodiversity and NCP risk. These areas will benefit by a reduction in anthropogenic pressure, e.g. through the establishment (or reinforcement) of protected areas and other area-based conservation measures (Nelson & Chomitz, 2011; Pennington et al., 2018). Existing protected areas might require expansion, to allow for shifting species ranges. The creation of corridors will also be required for species expected to face unsuitable climatic conditions, and with the ability to disperse to new areas. Areas with high risk of reduced regulation of water quantity (NCP 6) will additionally require the management of water supplies, reducing cropland expansion and deforestation which increase the runoff and flow speed. In the short term, changes in land-use management can promote water conservation through efficient agricultural policies, preventing water loss and agricultural runoff (IPBES, 2019; OECD Secretariat, 2015). Improved land-use management will also be essential for reducing the risk associated with wildfire (Lindenmayer et al., 2022; Ward et al., 2022).

In the recent years, there has been a rise in the number of disasters linked to extreme fires, impacting human population and wildlife in places such as Chile and Portugal in 2017, Greece in 2018, California in 2018 and 2020, and Australia in 2019 and 2020. These events have led to increased concern about the effect and range of wildfire, emphasizing the necessity to develop effective mitigation and adaptation strategies (Richardson et al., 2022). Extreme heatwaves and aridity add to this risk and generated potentially synergistic effects over extinction risk (Brook et al., 2008). These same threats affect some of the key contributions that nature delivers to people (Chaplin-Kramer et al., 2019, 2022; Clarke et al., 2022), generating high-risk areas for species and NCP. Bolder climate mitigation action will be globally needed to reduce the risk of triggering climatic tipping points (McKay et al., 2022). At the same time, climate adaptation will be needed in several areas even if global warming remains within Paris-set limits. We argue that intervention strategies that address areas facing both biodiversity and human risk, as identified here, should receive priority attention, as the effects of climate change will span multiple systems. Strategic planning and financial support of biodiversity conservation can deliver a high benefit to NCP preservation (Jung et al., 2021), and action to abate threats locally should be made rapidly (including sustainable land-use and water management) before global warming reduces the possibility to achieve win-win environmental solutions.

## Data availability

All input data used in these analyses derive from published sources cited in the methods section.

The CMIP6 datasets analysed during the current study are available from (https://esgf-node.llnl.gov/search/cmip6/). Computer code used in the analysis and outputs are openly available on Zenodo at https://zenodo.org/records/12799152.

