## Supplementary Material for "Areas of high risk for mammalian biodiversity and Nature’s Contributions to People under global warming"

### Table S1: List of the threats used for selecting species part of our analysis, from the IUCN Classification of Direct Threats.

| **Threat** | **Definition** |
| --- | --- |
| Fire & Fire Suppression (7.1) | Suppression or increase in fire frequency and/or intensity outside of its natural range of variation. |
| Increase in Fire Frequency/Intensity (7.1.1) | List the specific source of fire e.g., inappropriate fire management, escaped agricultural fires, arson, campfires, fires for hunting, etc |
| Droughts (11.2) | Periods in which rainfall falls below the normal range of variation |
| Temperature Extremes (11.3) | Periods in which temperatures exceed or go below the normal range of variation |

### **Table S2:** List of life-history traits and environmental variables considered as predictors of extinction risk in the random forest model before check for VIF. For the random forest model we kept only variables with VIF < 4. Data sources listed here have been cited in the main text.

| **Variable name** | **Definition** | **Unit of measures** | **Data source** |
| --- | --- | --- | --- |
| Adult mass | Body mass of an adult individual | grams | COMBINE |
| Gestation length | Length of time of foetal growth | days | COMBINE |
| Interbirth interval | Time between reproduction events | days | COMBINE |
| Consecutive dry days (cdd) | Maximum number of days in a row with precipitation below 1 mm in a year | days | ETCCDI |
| Warm days (TX90p) | Percentage of days with maximum temperature above the corresponding calendar day 90th percentile of maximum temperature for a 5-day moving window in the base period | % | ETCCDI |
| Minimum value of daily maximum temperature (TXn) | Minimum of daily maximum temperature in period (year or month). | °C | ETCCDI |
| Warm spell duration index (wsdi) | Annual count of days with at least 6 consecutive days when daily maximum temperature is above the calendar day 90th percentile of maximum temperature centered on a 5-day sliding window during the base period | days | ETCCDI |
| Length of the fire season (fwils) |  | days | FWI |
| Seasonal average of the FWI (fwisa) |  | days | FWI |
| Number of days with extreme fire weather (fwixd) |  | days | FWI |
| Maximum value of the FWI (fwixx) |  | Integer | FWI |
| Maximum Actual Evapotranspiration (AET max) | Maximum annual actual evapotranspiration | mm/month | CMIP6 |
| Minimum Actual Evapotranspiration (AET min) | Minimum annual actual evapotranspiration | mm/month | CMIP6 |
| Mean Actual Evapotranspiration (AET mean) | Mean annual actual evapotranspiration | mm/month | CMIP6 |
| Minimum Water Availability (WA min) | Minimum annual water availability | mm/month | CHELSA /CMIP6 |
| Maximum Water Availability (WA max) | Maximum annual water availability | mm/month | CHELSA /CMIP6 |
| Mean Water Availability (WA avg) | Mean annual water availability | mm/month | CHELSA /CMIP6 |
| Maximum Burned Area | Maximum annual burned area | kg/m^2^ | CMIP6 |
| Minimum Burned Area | Minimum annual burned area | kg/m^2^ | CMIP6 |
| Mean Burned Area | Mean annual burned area | kg/m^2^ | CMIP6 |
| Maximum CO2 Emission | Maximum annual CO2 Emission from fire | kg/m^2^ | CMIP6 |
| Minimum CO2 Emission | Minimum annual CO2 Emission from fire | kg/m^2^ | CMIP6 |
| Mean CO2 Emission | Mean annual CO2 Emission from fire | kg/m^2^ | CMIP6 |
| Anthropic | Sum of C3 annual crops, C3 perennial crops, C3 nitrogen-fixing crops, C4 annual crops and C4 perennial crops | km^2^ | LUH |

### Table S3: Table reporting the GCMs available for each NCP indicator.

| NCP  GCM | Precipitation | Actual Evapotranspiration | Water Availability (NCP6) | Burned area (NCP9) | CO_2_ emissions from fire (NCP3) |
| --- | --- | --- | --- | --- | --- |
| UKESM1-0-LL | 1 | 1 | 1 | 0 | 0 |
| MRI-ESM2-0 | 1 | 1 | 1 | 0 | 0 |
| IPSL-CM6A-LR | 1 | 1 | 1 | 0 | 0 |
| MPI-ESM1-2-HR | 1 | 1 | 1 | 0 | 0 |
| GFDL-ESM4 | 1 | 1 | 1 | 0 | 1 |
| CESM2-WACCM | 0 | 0 | 0 | 1 | 1 |
| CMCC-ESM2 | 0 | 0 | 0 | 1 | 1 |
| EC-Earth3-Veg-LR | 0 | 0 | 0 | 0 | 1 |
| NorESM2-LM | 0 | 0 | 0 | 1 | 1 |
| NorESM2-MM | 0 | 0 | 0 | 1 | 1 |
| MPI-ESM1-2-LR | 0 | 0 | 0 | 0 | 1 |
| CMCC-CM2-SR5 | 0 | 0 | 0 | 1 | 0 |

### Table S4: Results of the taxonomic cross-validation (family level, number of species >15) for the random forest models in terms of: Sensitivity, Specificity, True Skills Statistics (TSS).

| **Family** | **n °species in the family** | **Sensitivity** | **Specificity** | **TSS** |
| --- | --- | --- | --- | --- |
| Cricetidae | 15 | 0.67 | 0.75 | 0.42 |
| Muridae | 36 | 0.59 | 0.58 | 0.17 |
| Bovidae | 29 | 0.41 | 0.42 | -0.17 |
| Sciuridae | 15 | 0.57 | 0.62 | 0.2 |
| Pteropodidae | 17 | 0.71 | 0.7 | 0.41 |
| Dasyuridae | 16 | 0.5 | 0.5 | 0 |
| Vespertilionidae | 36 | 0.56 | 0.56 | 0.11 |
| Leporidae | 16 | 0.62 | 0.62 | 0.25 |
| Nesomyidae | 18 | 0.6 | 0.69 | 0.29 |
| Carcopithecidae | 21 | 0.67 | 0.67 | 0.33 |
| Soricidae | 15 | 0.83 | 0.78 | 0.61 |
| Macropodidae | 19 | 0.62 | 0.64 | 0.26 |
| Tenrecidae | 16 | 0.67 | 0.77 | 0.44 |

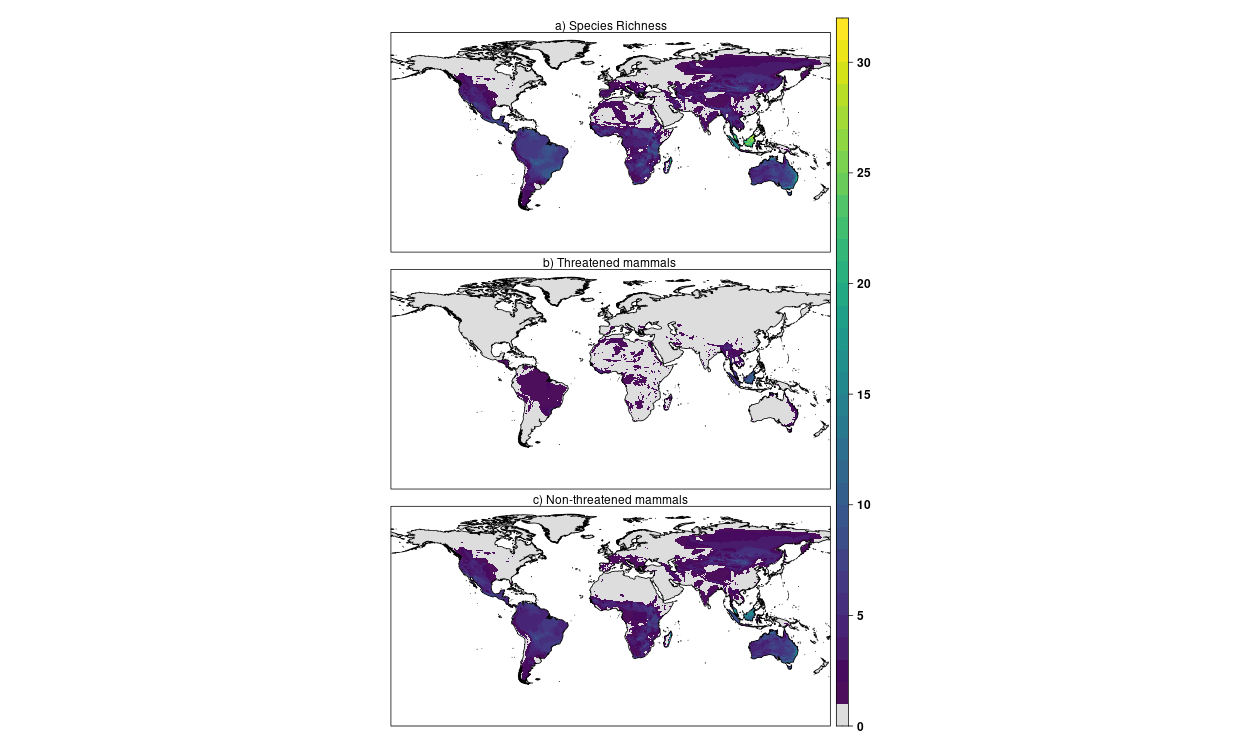

### **Fig. S1:** Map representing a) the number of mammal species in our analysis (colorbar) for each 10km grid cell of the globe; b) the number of threatened mammal species in our analysis (colorbar) for each 10km grid cell of the globe. c) the number of non-threatened mammal species in our analysis (colorbar) for each 10km grid cell of the globe.

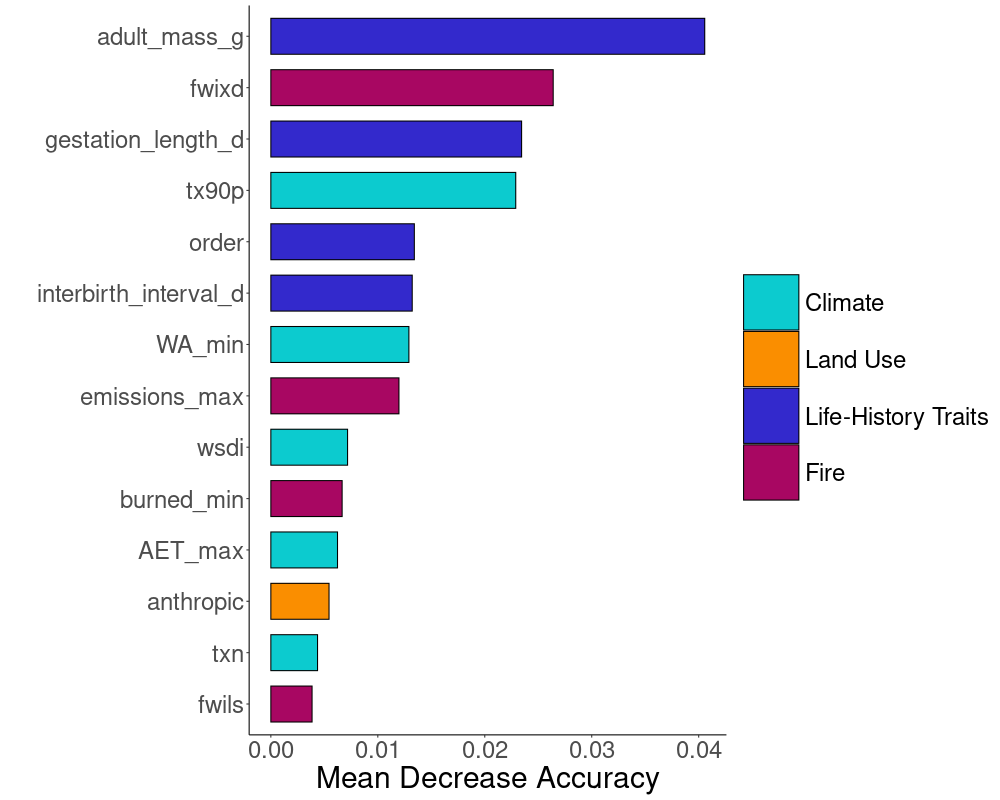

### Fig. S2: Random Forest variable importance plot for selected predictors in the final extinction risk model.

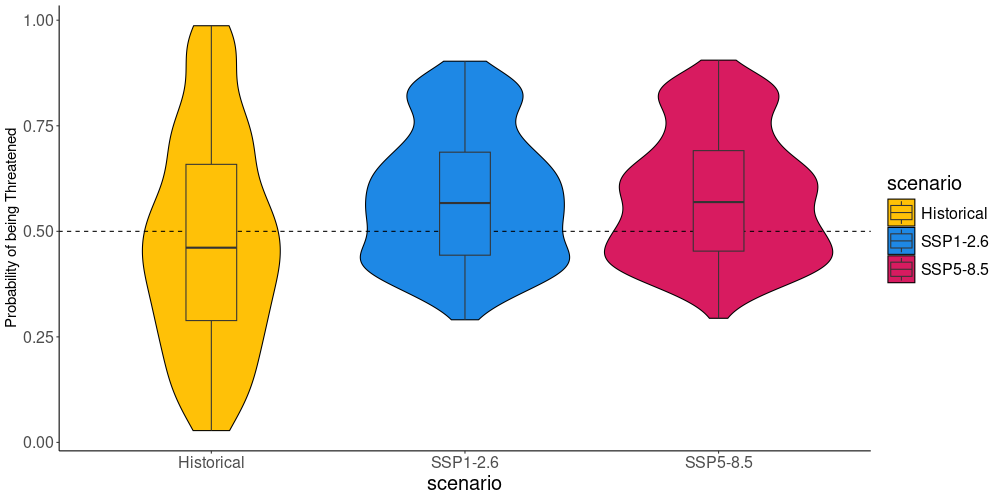

### Fig. S3: Box plot showing the probability of being threatened for species under the historical time period (1981-2010) the two scenarios of climate change (2041-2070): SSP1-2.6 and SSP5-8.5.

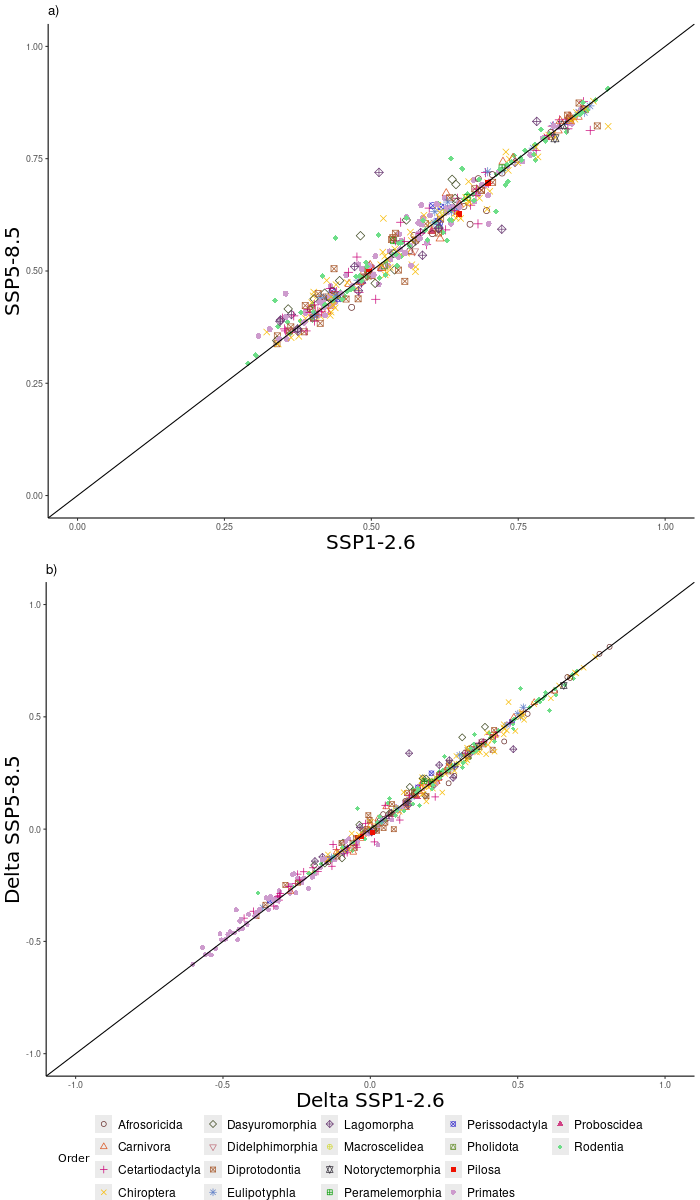

### Fig. S4: Scatter plot showing a) the probability of being threatened and b) the change in the probability of being threatened for species under the two scenarios of climate change (2041-2070): SSP1-2.6 and SSP5-8.5. Species are coloured by the order they belong to.

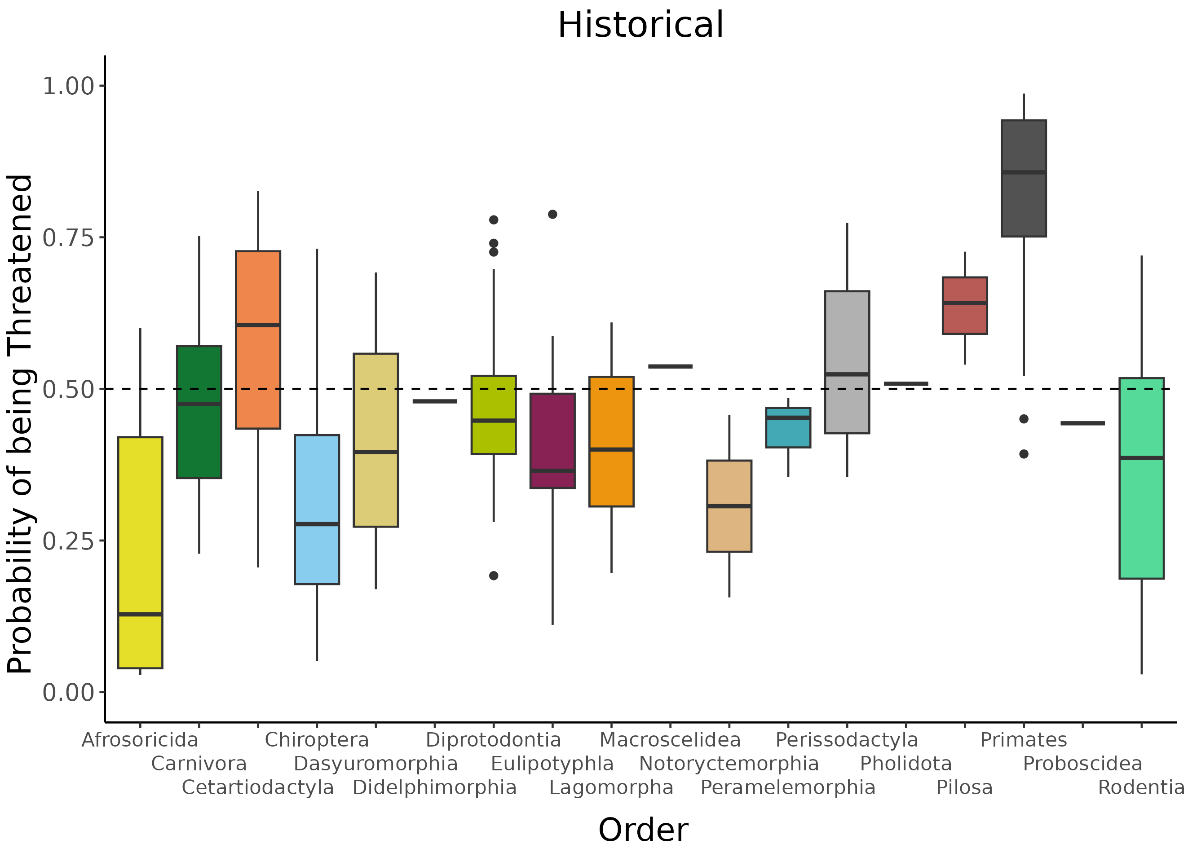

### Fig. S5: Probability of being threatened for species in different taxonomic orders, under historical period 1981-2010.

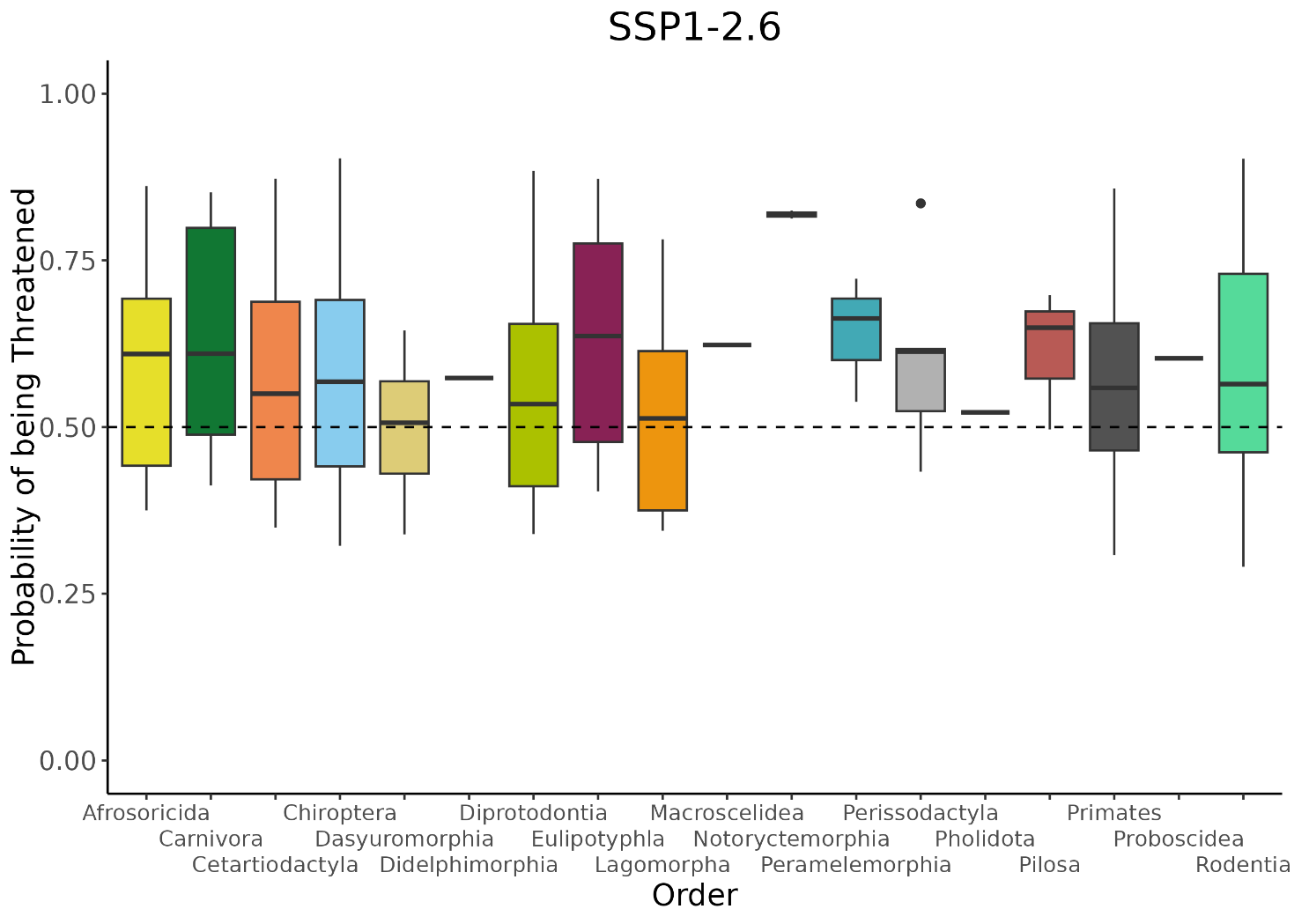

### Fig. S6: Probability of being threatened for species in different taxonomic orders, under the scenario SSP1-2.6.

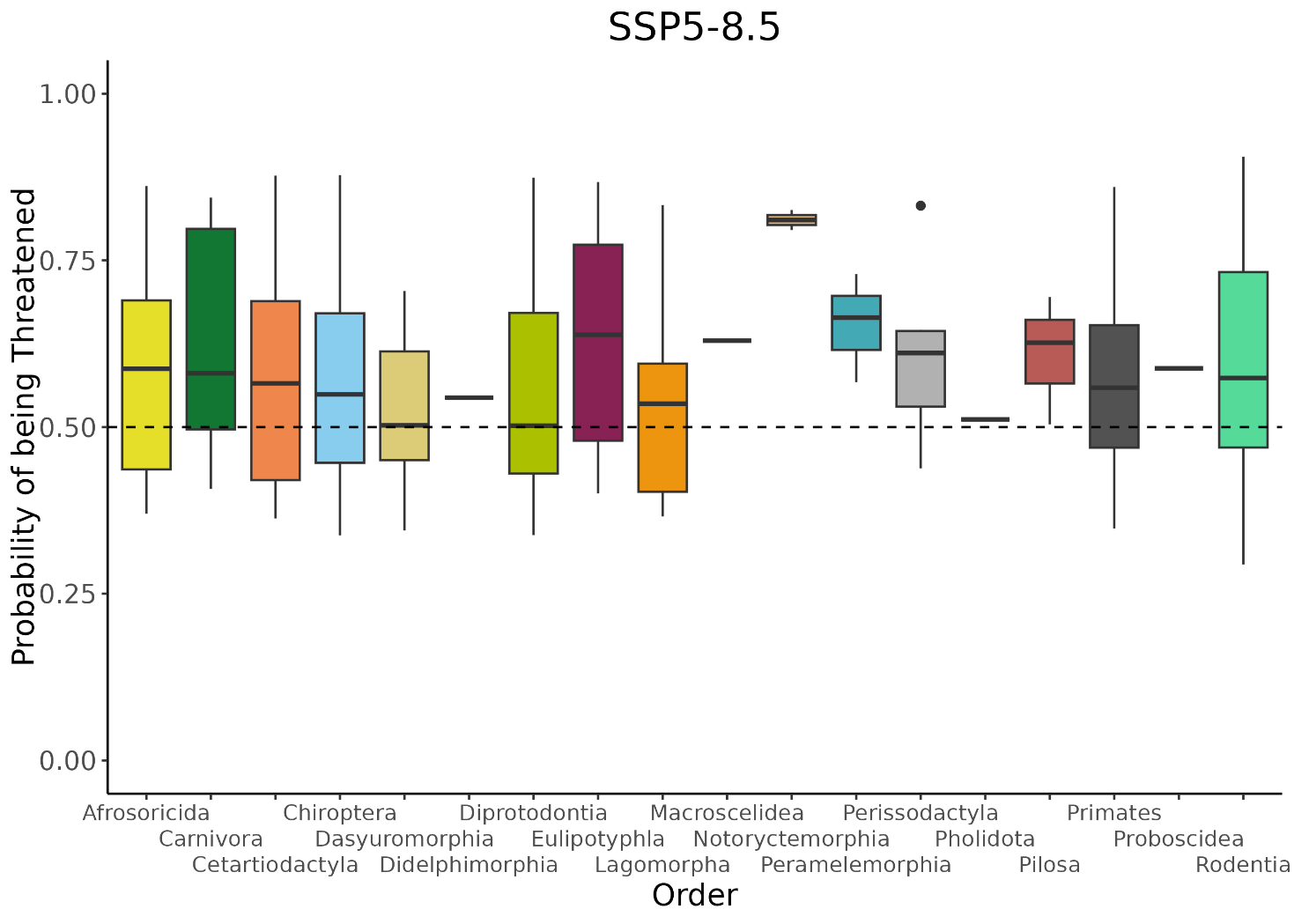

### Fig. S7: Probability of being threatened for species in different taxonomic orders, under the scenario SSP5-8.5.

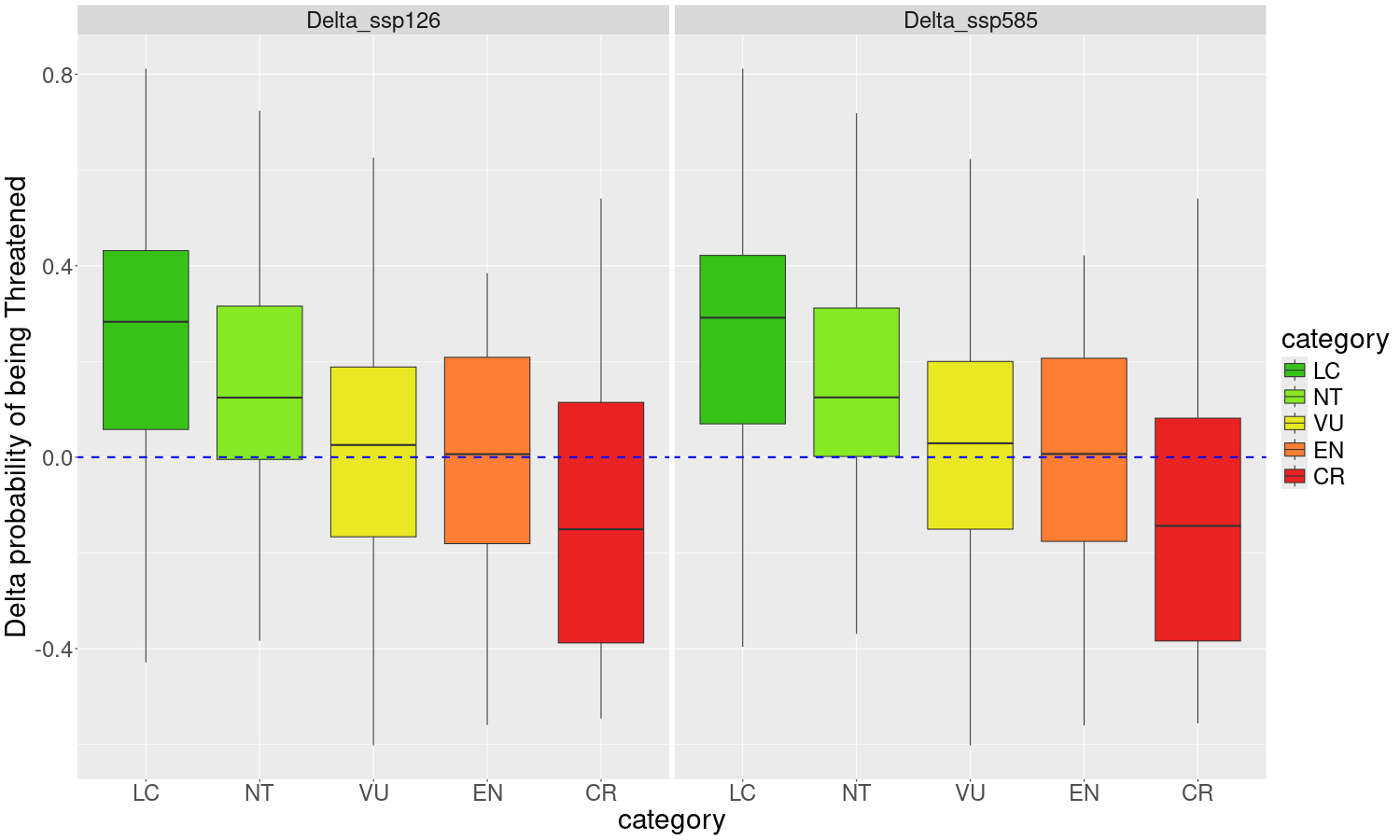

### **Fig. S8:** Change in the probability of being threatened for species in different IUCN Red List Categories, under the two scenarios of future climate change SSP1-2.6 and SSP5-8.5. Categories are as follow: CR = Critically Endangered; EN = Endangered; VU = Vulnerable; NT = Near Threatened; LC = Least Concern.

**.**
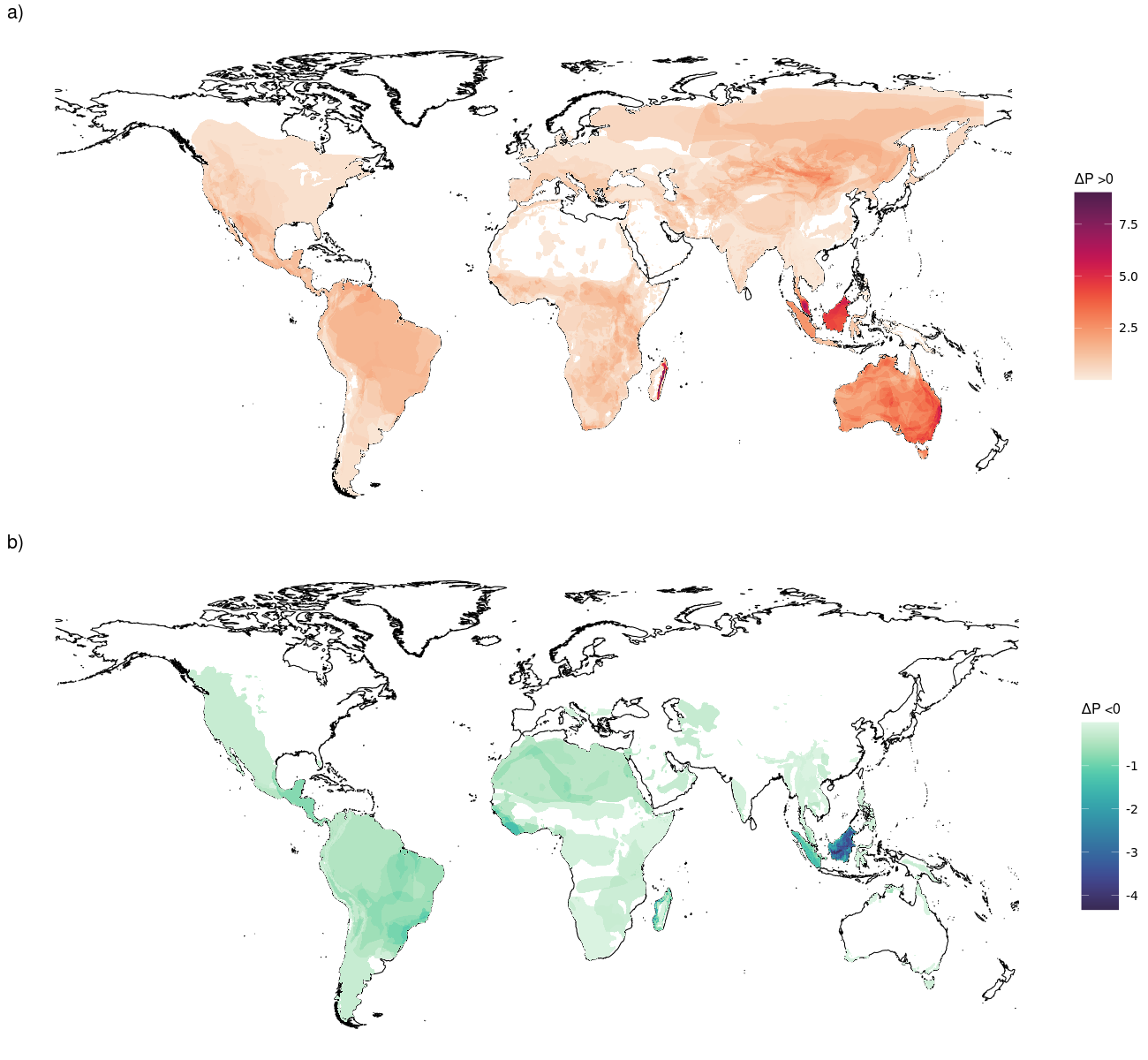

### **Fig. S9**: Global cumulative change in species extinction risk for species showing a) a risk increase and b) a risk decrease, under scenario SSP1-2.6.

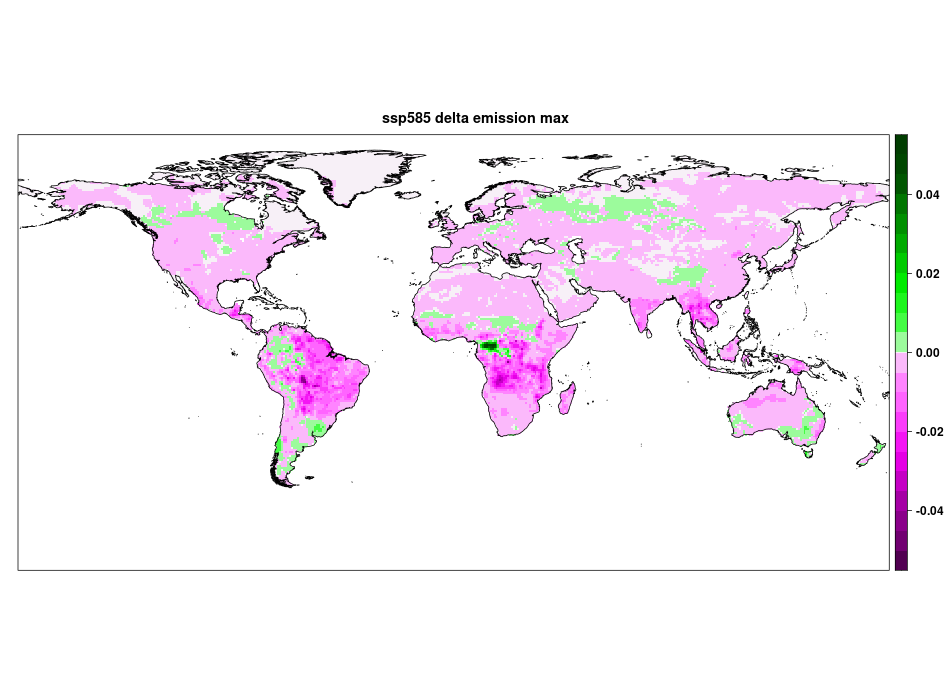

### **Fig. S10:** Projections of change in the value of maximum CO2 emission from fire between the baseline period 1981-2010 and the future 2041-2070, under scenario SSP5-8.5.

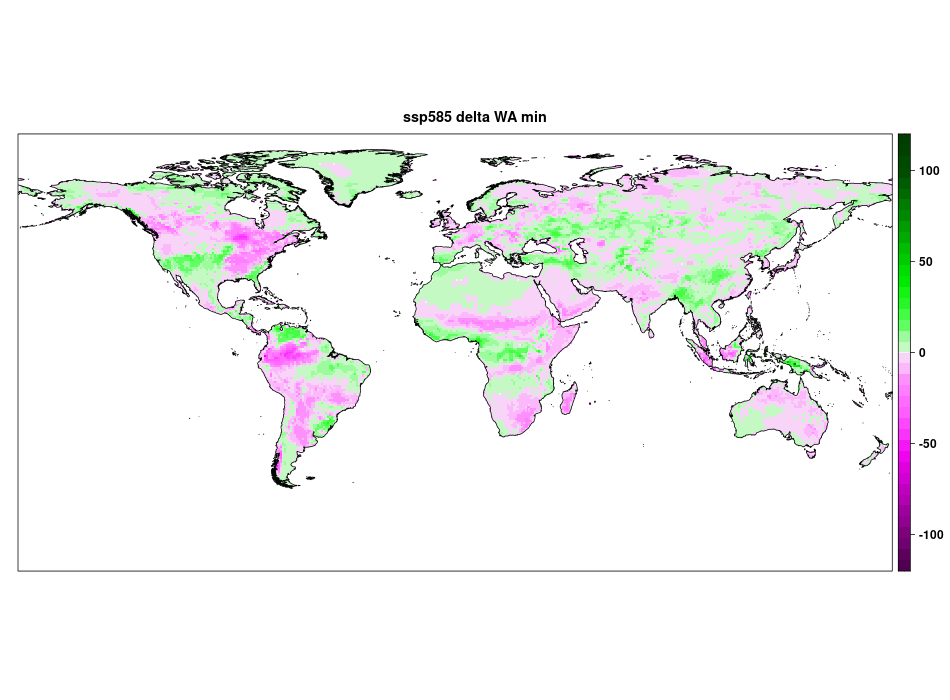

### **Fig. S11:** Projections of change in the value of minimum WA between the baseline period 1981-2010 and the future 2041-2070, under scenario SSP5-8.5.

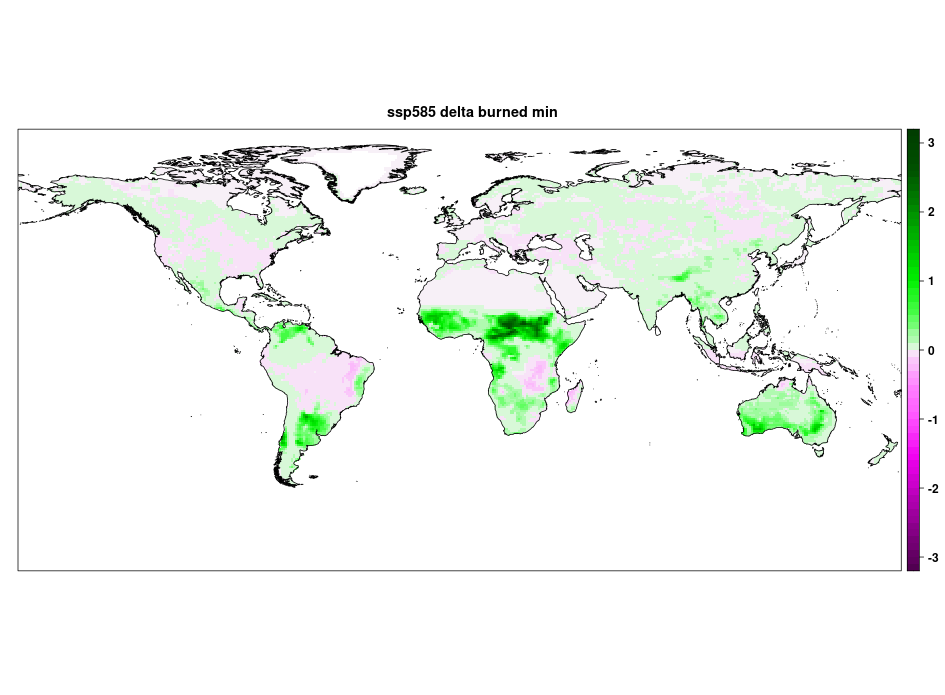

### **Fig. S12:** Projections of change in the value of minimum burned area between the baseline period 1981-2010 and the future 2041-2070, under scenario SSP5-8.5.

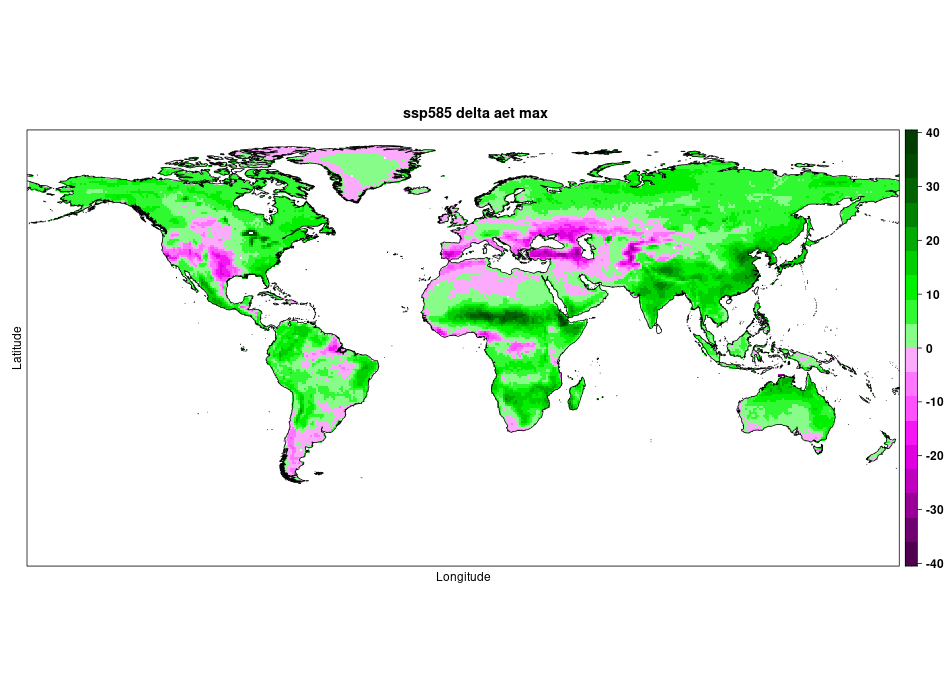

### **Fig. S13:** Projections of change in the value of maximum actual evapotranspiration between the baseline period 1981-2010 and the future 2041-2070, under scenario SSP5-8.5.

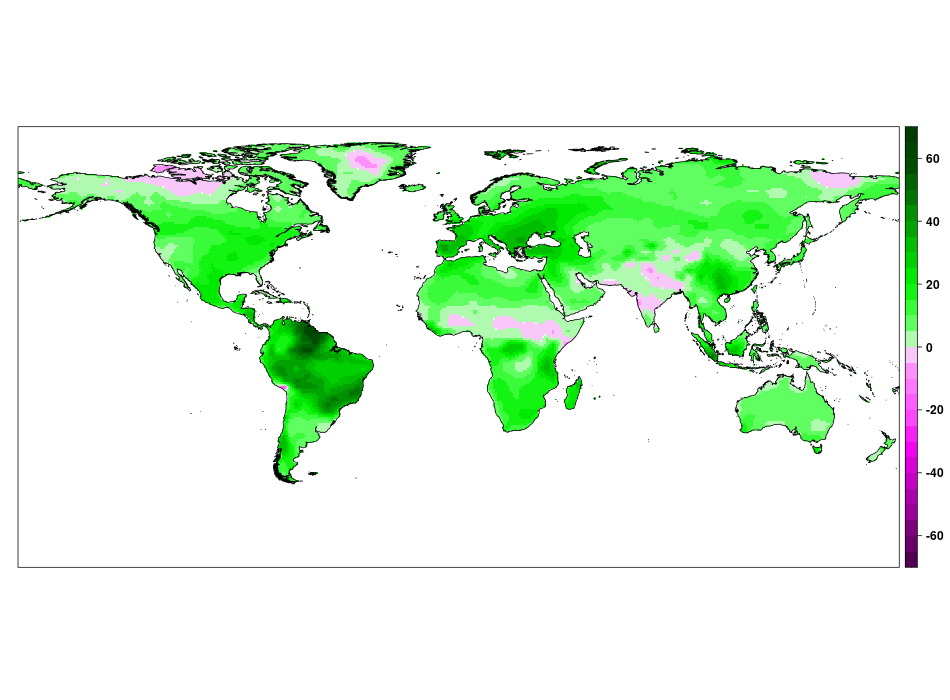

### Fig. S14: Projections of change in the value of number of days with extreme fire weather between the baseline period 1981-2010 and the future 2041-2070, under scenario SSP5-8.5.

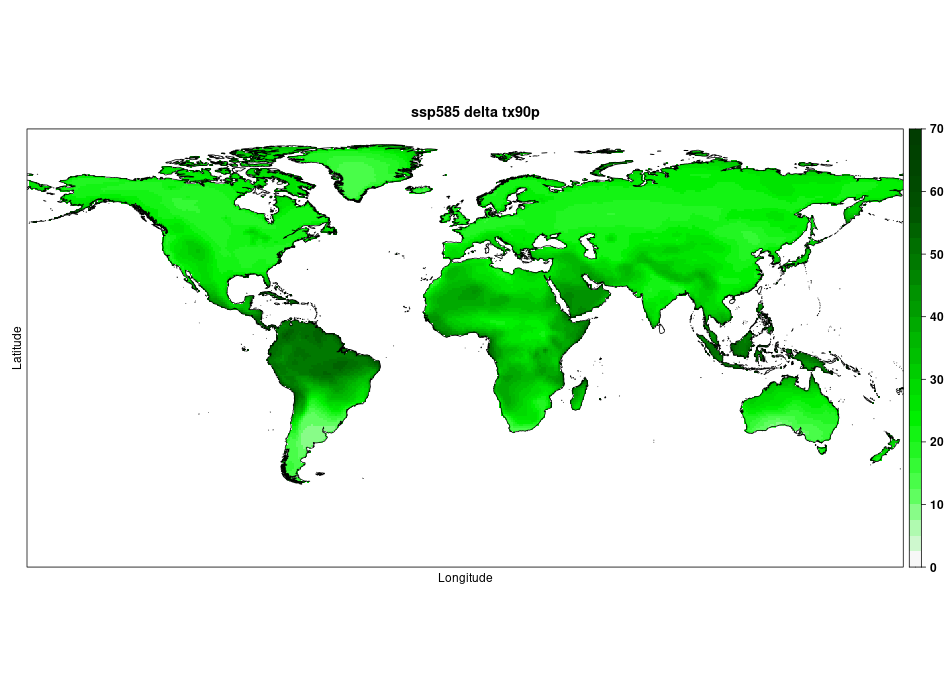

### Fig. S15: Projections of change in the value of warm days between the baseline period 1981-2010 and the future 2041-2070, under scenario SSP5-8.5. Warms days are defined as the percentage of days with maximum temperature above the corresponding calendar day 90th percentile of maximum temperature for a 5-day moving window in the base period.

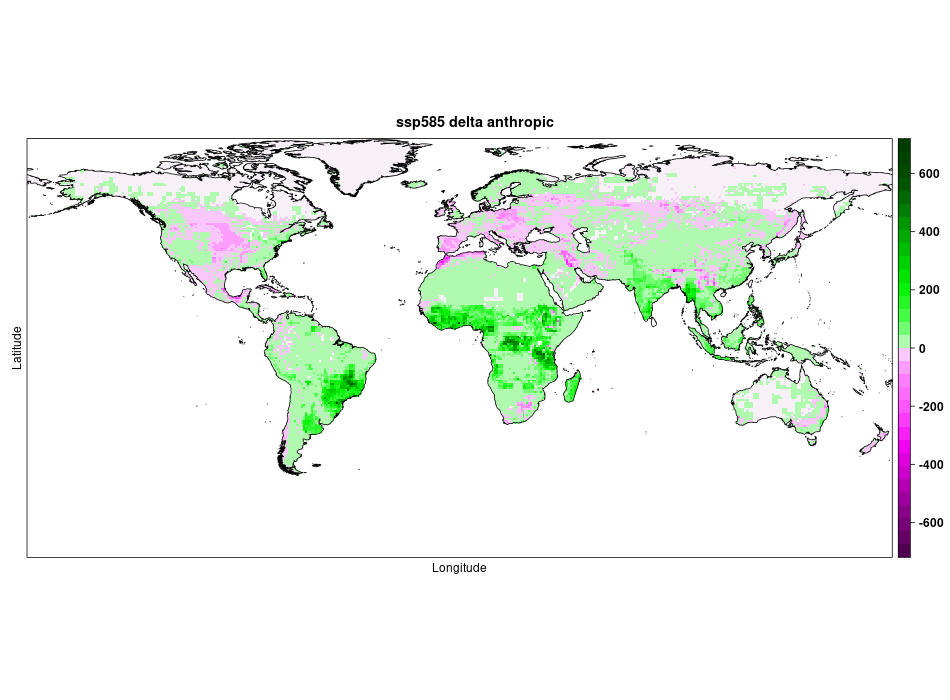

### Fig. S16: Projections of change in the value of anthropic land between the baseline period 1981-2010 and the future 2041-2070, under scenario SSP5-8.5.

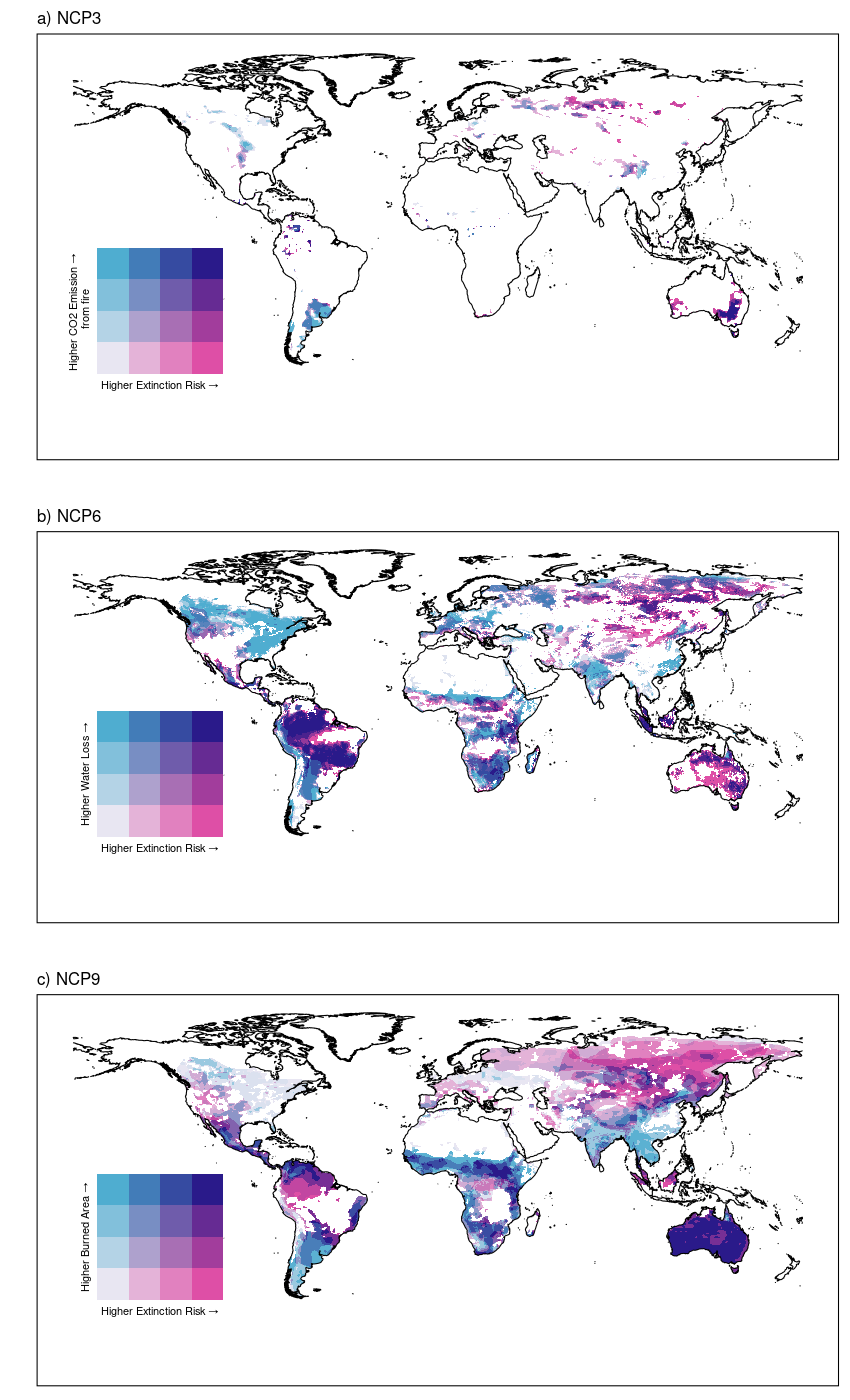

### **Fig. S17:** Global bivariate maps of increase in species extinction risk versus NCP indicators, specifically CO_2_ fire emission, indicator of Air quality regulation (NCP 3), water availability indicator of freshwater quantity regulation (NCP 6) and Burned area, indicator of regulation of extreme event (NCP 9). The map reports relative increase in extinction risk for species expected to have higher risk in the future and increase the proxy for NCP under scenario SSP1-2.6. High-risk area of species and NCP risk are identified in purple.

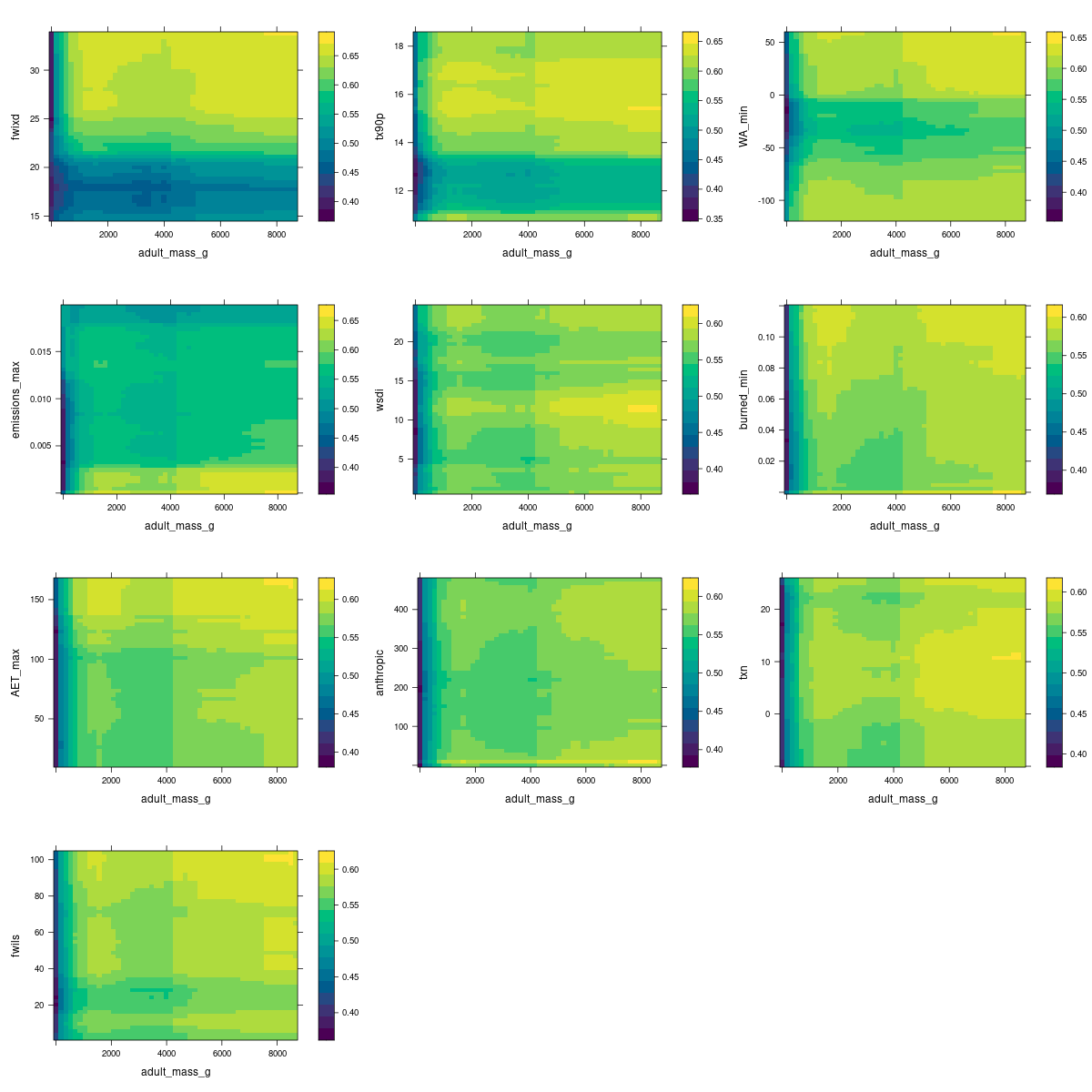

### Fig. S18: Bivariate partial dependency plots showing response of the probability of being threatened to adult mass vs extrinsic variables (yellow=higher probability of being threatened). In order, extreme fire weather days (fwixd), warm days (tx90p), minimum water availability (WA_min), maximum CO2 fire emission (emission_max), warm spell duration index (wsdi), minimum burned area (burned_min), maximum actual evapotranspiration (AET_max), anthropic land (anthropic), Minimum of daily maximum temperature(txn) and length of fire season (fwils) .
